# Acute Myeloid Leukemia Relapse after Bromodomain Inhibitor Treatment or Chemotherapy is Characterized by Myc-Ras Transcriptional Remodeling

**DOI:** 10.1101/2025.11.13.688312

**Authors:** Benjamin J. Huang, Jeevitha D’Souza, Alexa Rane Batingana, Max D. Harris, Xinyue Wang, Eugene Hwang, Anica M. Wandler, Michael R. Burgess, Qing Li, Soheil Meshinchi, Gideon Bollag, Kevin Shannon

## Abstract

Adult and pediatric acute myeloid leukemias (AMLs) harbor distinct mutational profiles, including a higher incidence of *RAS* and other signaling mutations in young patients. Here we show that the BET inhibitor PLX51107 potently suppresses the growth of *NRAS*-mutant AML cell lines, and that these activities are enhanced by co-treatment with the MEK inhibitor PD0325901. Controlled preclinical trials in primary mouse *Nras*-mutant AMLs revealed single agent efficacy of PLX51107 that was enhanced by PD0325901. Leukemias that relapsed during treatment developed intrinsic drug resistance characterized by transition to a more primitive state, up-regulation of Myc target genes, and down-regulation of Ras-associated transcriptional programs. AMLs that relapsed after frontline chemotherapy showed similar transcriptional remodeling. These studies demonstrate transcriptional plasticity in primary AMLs that relapse following *in vivo* treatment with either targeted agents or chemotherapy, and support evaluating BET inhibition in leukemias with monocytic differentiation and *RAS* mutations.

## INTRODUCTION

Acute myeloid leukemia (AML) is an aggressive and molecularly heterogeneous cancer that affects all age groups ^1^. Despite recent progress in developing new agents, front-line therapeutic protocols have not changed markedly over the past two decades and are associated with substantial morbidity and mortality. Accordingly, there is a need to develop mechanism-based treatments and more effective drug combinations.

Somatic *NRAS, KRAS,* and *NF1* mutations occur in ∼20% of adult and in >50% of pediatric AMLs profiled at diagnosis ^1^. These and other signaling mutations track closely with disease activity in AML as they become undetectable during remission and reappear or are replaced by a different signaling mutation at relapse ^1–4^. In addition to playing a key role in AML pathogenesis, *NRAS* mutations are a major cause of *de novo* or adaptive resistance to the FLT3 inhibitor gilteritinib ^4^, the IDH2 inhibitor enasidenib ^5^, and the Bcl-2 inhibitor venetoclax ^6–8^.

Most oncogenic *RAS* mutations encode missense substitutions at codons 12, 13, and 61 that result in constitutively elevated levels of Ras-GTP due to reduced intrinsic GTP hydrolysis and resistance to GTPase activating proteins (GAPs). The mitogen activated protein kinase (MAPK) effector pathway is a key downstream target of Ras-GTP, and chemical inhibitors of Raf and MEK are approved by the Food and Drug Administration for the treatment of melanoma and other cancers. However, allosteric MEK inhibitors such as trametinib and selumetinib have shown disappointing efficacy as single agents in AML and most other advanced cancers ^9,10^.

The recent development of the K-Ras^G12C^ inhibitors adagrasib and sotorasib demonstrated the feasibility of directly targeting oncogenic Ras proteins and showed that primary *KRAS*-mutant cancers with multiple cooperating mutations are addicted to hyperactive Ras signaling *in vivo* ^11^. Similarly, genetic knockdown of oncogenic *NRAS*/*Nras* inhibits the growth of human AML cell lines *ex vivo* and of primary mouse leukemias *in vivo* ^12^. Recent studies of patients with lung adenocarcinoma who progressed after receiving K-Ras^G12C^ inhibitors revealed selective pressure for tumors to restore oncogenic Ras signaling through both on- and off-target adaptive resistance mechanisms ^13,14^. These clinical observations are consistent with previous work that emphasized the importance of developing effective strategies to potently inhibit mutant Ras signal output as well as targeting possible bypass mechanisms and co-existing mutations ^15,16^.

BET proteins are epigenetic “readers” that bind histones at acetylated lysine residues, such as acetylated H3K27, to recruit transcriptional regulatory complexes and promote the expression of nearby genes ^17^. Studies in a mouse model in which hematopoietic stem and progenitor cells (HSPCs) were transformed *ex vivo* by *KMT2A::AF9* (previously known as *MLL::AF9*), support the hypothesis that loss of polycomb repressor complex (PRC2) function promotes an epigenetic state that is both permissive for AML development and essential for leukemia maintenance ^18^. Broadly speaking, PRC2 antagonizes BET-related transcriptional activation by tri-methylating H3K27, which results in transcriptional silencing. Zuber *et al.* generated a transplantable AML model by co-expressing a KMT2A-AF9 fusion protein and N-Ras^G12D^ in mouse HSPCs ^19^ and used this system to perform an RNA interference screen that identified the BET protein Brd4 as essential for AML maintenance *in vivo* ^20^. Although the “first generation” BET inhibitor JQ1 showed anti-leukemia activity in this model, another compound (OTX015) had only modest efficacy in adult patients with relapsed AML ^21^. A recent trial that combined the BET inhibitor PLX51107 and the hypomethylating agent azacitidine reported an objective response rate of 22% in a group of heavily pretreated adult patients ^22^, suggesting improved combinations are warranted. Furthermore, a different BET inhibitor (pelabresib) and the JAK2 inhibitor ruxolitinib recently showed promising efficacy in patients with treatment-naïve myelofibrosis ^23^.

This study was informed by elegant mechanistic and preclinical data that revealed potent effects of combining JQ1 and ruxolitinib in a *Mpl^V515L^* mouse model of myelofibrosis ^24^.

Recent studies have provided evidence that AML differentiation state modulates drug response in some contexts ^25^. For example, Pei, et al. showed that primitive AML subtypes are more responsive to venetoclax, and that subsequent monocytic differentiation is associated with adaptive resistance ^26,27^. Furthermore, a comprehensive analysis of *ex vivo* drug responses within the BEAT AML cohort suggests that the degree of myeloid differentiation broadly impacts drug sensitivity. Specifically, these investigators showed that AMLs with a more differentiated transcriptional state were sensitive to trametinib, selumetinib, and JQ1 ^25^. These data are consistent with previous studies showing that *RAS* mutations and *KMT2A*-rearrangements are enriched in more differentiated AML morphological subtypes ^28^. Altogether, these findings provide a strong rationale for evaluating the efficacy of combined BET and MEK inhibition in AMLs characterized by arrested monocytic differentiation, particularly leukemias that harbor *RAS* mutations.

PLX51107 is well-tolerated after *in vivo* administrations at a 10-fold lower dose than OTX015, suggesting a broader therapeutic index ^29^. Here we show that PLX51107 is highly synergistic with the MEK inhibitor PD0325901 (PD901) in *NRAS/Nras*-mutant, monocytic AML cell lines and murine AML models. Primary leukemias that relapse during continuous treatment with PLX51107 or with the PLX51107/PD901 combination develop intrinsic drug resistance that is associated with a transition to a more primitive differentiation state. This transition is characterized by down-regulation of Ras-associated transcriptional programs and up-regulation of Myc-regulated gene sets and is independent of Myc protein expression levels. Bulk and single cell transcriptome analysis from pediatric AML patients treated with frontline cytotoxic chemotherapy uncovered a similar fate shift whereby leukemias transition towards stem-like transcriptional signatures at relapse while simultaneously up- and down-regulating Myc- and Ras-associated programs, respectively. Altogether, these studies indicate that remodeling of transcriptional networks associated with hematopoietic differentiation is a common feature of adaptive resistance in AMLs treated with BET inhibitors or standard-of-care chemotherapy.

Furthermore, our data support evaluating combined BET and MEK inhibition in *NRAS*-mutant, monocytic subtypes of AML. PLX51107 development is continuing at Opna Bio LLC as OPN-51107.

## RESULTS

### *RAS* Mutations and Ras-Regulated Gene Signatures Are Associated with Myeloid Differentiation in Pediatric AML

Sensitivity to MEK inhibition is associated with more differentiated AML transcriptional states in adult AML ^25^. To assess potential relationships between *RAS* mutations, Ras-regulated transcriptional programs, and differentiation states in pediatric AML, we performed gene expression analysis and variant calling on a large whole transcriptome RNA sequencing (RNA-seq) dataset generated as part of the NCI TARGET Initiative (N = 760) (**Fig. 1A**). Starting with the FAB (French-American-British) morphologic classification of AML (**Fig. 1B**), we asked whether myeloid differentiation is associated with a higher prevalence of *NRAS* and *KRAS* mutations. Consistent with previous findings ^28^, the frequency of *RAS* mutations was highest in AML subtypes characterized by myelomonocytic or monocytic maturation (FAB M4/M5) (**Fig. 1C**). Next, we confirmed that FAB morphologic classification is closely associated with transcriptional signatures of myeloid maturation (**Fig. 1D**). The degree of myeloid maturation was uniquely correlated with up-regulation of *RAS* gene signatures (R^2^ = 0.8558 and 0.8467), but not other signal transduction or survival programs (**Fig. 1E, 1F, and Fig. S1**). Intriguingly, this association of morphologic and transcriptional differentiation with *RAS* gene signatures was independent of *RAS* mutation status.

**Figure 1.**
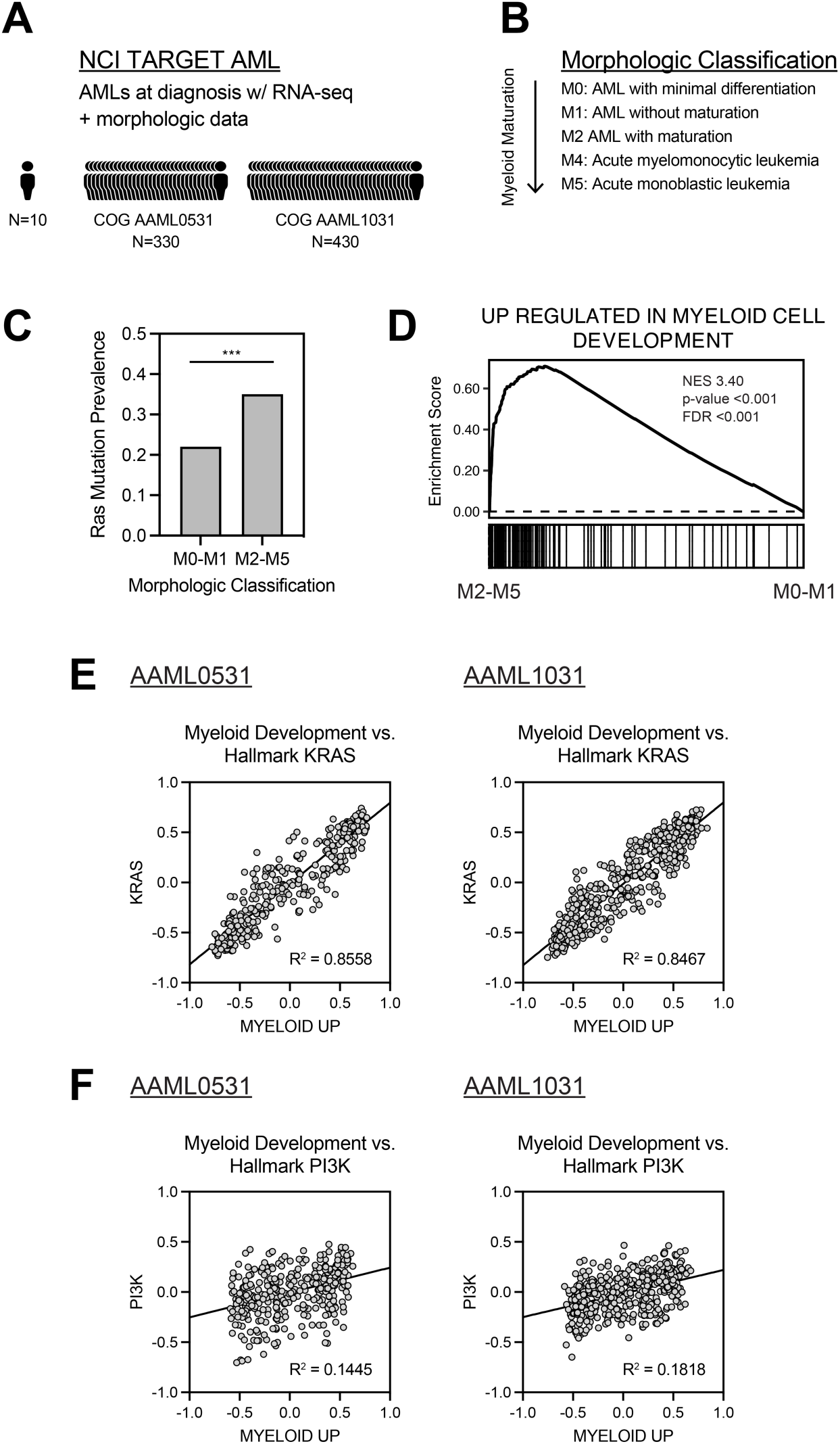
Morphologic maturation and myeloid differentiation gene sets are associated with increased Ras transcriptional output. **A.** NCI TARGET AML transcriptome data (N = 760) were analyzed. Bulk RNA sequencing (RNA-seq) and morphology data were available from 330 patients enrolled on Children’s Oncology Group (COG) clinical trial AAML0531 and from 430 patients enrolled on COG clinical trial AAML1031. **B.** Categorization was based on AML French-American-British (FAB) morphologic classification where increasing differentiation is seen from M0 to M5. **C**. *RAS* mutations are more prevalent in FAB morphologic AML subtypes M2-M5 (compared to M0-M1 subtypes). **D.** A normal myeloid development transcriptional program is enriched in M2, M4, and M5 AMLs compared to M0 and M1 AMLs. **E, F.** GSVA enrichment scores for Broad Molecular Signature Database myeloid development versus Ras signature gene sets are highly correlated in AAML0531 and AAML1031 patient cohorts. By contrast, no relationship is observed for PI3K or other hallmark gene sets. See also **Figure S1**.

### BET and MEK Inhibition Synergistically Reduce Growth and Promote Apoptosis in *NRAS* Mutant AML Cell Lines

PLX51107 is a BET bromodomain inhibitor that potently inhibits the growth of most hematologic cancer cell lines at IC50 values in the nanomolar range, including six AML lines with a mutation in either *NRAS* or *KRAS* (**Table S1** and **Fig. 2A**) ^29^. Informed by our analysis of pediatric AML transcriptomes (**Fig. 1**) and *in vitro* data suggesting that monocytic AML subtypes may be sensitive to BET and MEK inhibitor combinations ^25^, we exposed the *NRAS*-mutant human monocytic AML cell lines OCI-AML3, HL-60, and THP-1 to varying and overlapping concentrations of PLX51107 and PD901. Experiments that utilized the CellTiter-Glo (CTG) assay as an endpoint measure of viable, metabolically active cells showed that PLX51107 and PD901 are highly synergistic based on log-fold shifts in dose response curves (**Fig. 2B and S2A**), isobologram analysis (**Fig. 2C**) and, Bliss independence scores (**Fig. 2D**). Synergistic activity was observed across a range of effective concentrations (EC90, EC75, EC50) (**Fig. 2C**) and a matrix of varying dose levels (**Fig. 2D**).

**Figure 2.**
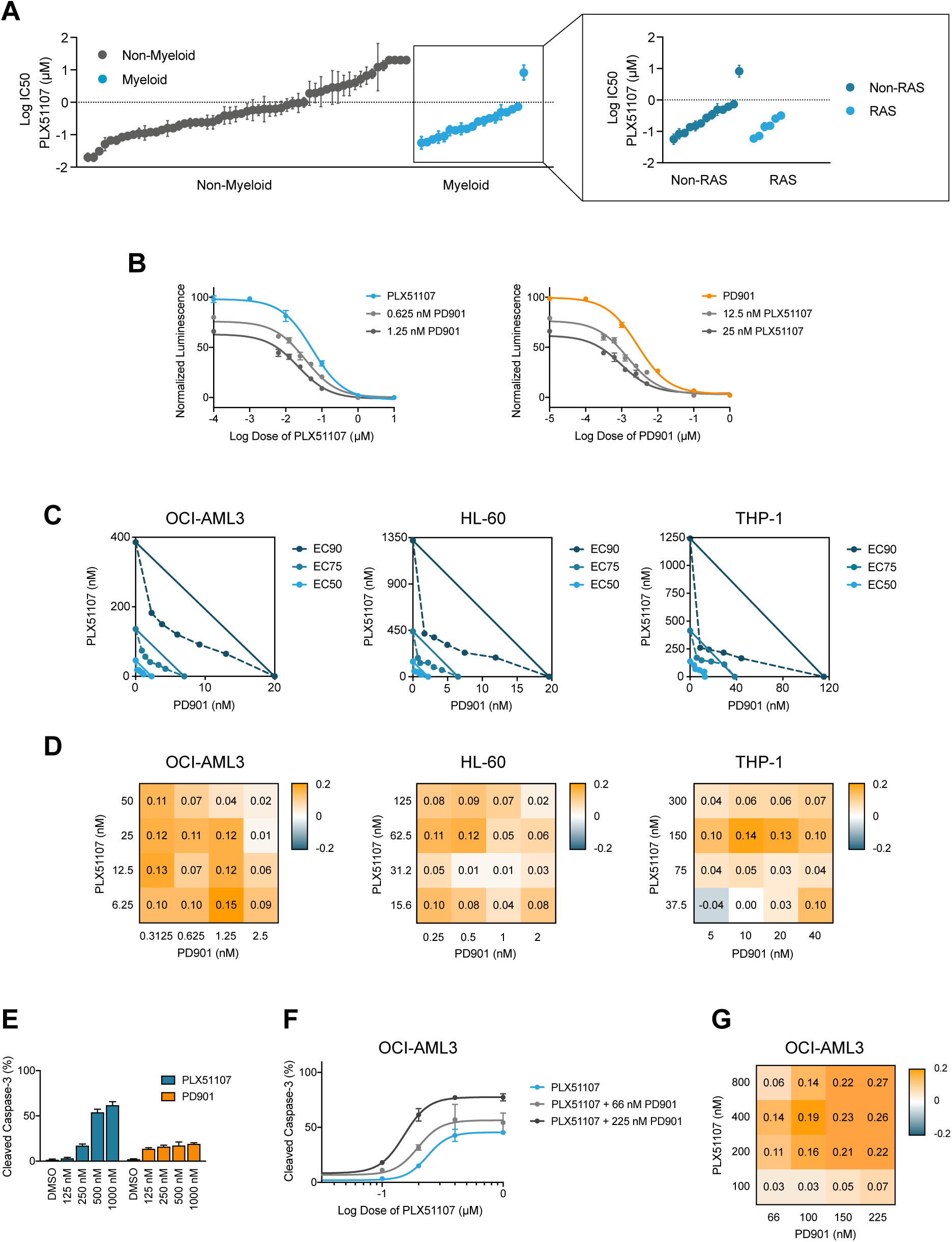
Activity of PLX51107 alone and in combination with PD901 in *NRAS*-mutant AML cell lines. **A.** Log10 IC50 for PLX51107 from a previous analysis of a panel of hematologic cancer cell lines ^29^ but with a specific focus on myeloid and *RAS*-mutant cell lines, which are uniformly sensitive to PLX51107 in nanomolar ranges. **B.** CellTiter-Glo (CTG) analysis of the *NRAS*-mutant cell line OCI-AML3 with single agent PLX51107 (blue) or PD901 (orange) and in combination at low nanomolar dose ranges (gray). **C.** Isobologram analysis of *NRAS*-mutant OCI-AML3, HL-60, and THP-1 AML cells shows synergy at 90%, 75%, and 50% effective concentration levels. **D.** Positive synergistic Bliss independence scores were observed across a range of PLX51107 and PD901 doses. **E.** PLX51107 induces apoptosis at high nanomolar doses in the OCI-AML3 cell line as measured be the percentage of cells expressing cleaved caspase-3, whereas PD901 is less effective at inducing apoptosis. **F.** Induction of apoptosis by PLX51107 is potentiated by PD901 at every dose level and is associated with a significant increase in maximum effect. **G.** Positive (synergistic) Bliss independence scores for CC3 staining were observed across a range of PLX51107 and PD901 doses. See also **Figure S2**.

Reduced growth as assessed by CTG analysis can result from cell death and/or proliferative arrest of viable cells. To distinguish between these two fates, we performed additional single agent and synergy testing utilizing cleaved caspase 3 (CC3) detection as a readout of apoptosis. Consistent with previous observations in primary *Nras* mutant mouse AML cells ^12^, PD901 treatment induced cell cycle arrest in human AML cell lines without substantially increasing the percentage of CC3-positive (CC3+) cells (**Fig. 2E**). By contrast, PLX51107 caused a significant and dose-dependent increase in cell death (**Fig. 2E**). Exposing OCI-AML3, HL-60, and THP-1 cells to varying and overlapping concentrations of PLX51107 and PD901 revealed potent synergy and increases in the maximum apoptotic effect at nanomolar concentrations of both drugs (**Fig. 2F and S2B**). Dose-response curves generated using CC3 staining did not share the same uniform maximum effect detected in CTG assays. While this precluded performing isobologram analysis, Bliss independence scores were robustly positive across several dose levels (**Fig. 2G and S2C**). By contrast, we observed no evidence of synergy between PLX51107 and PD901 in the chronic myeloid leukemia cell line K562, in which Ras is activated downstream of the BCR-ABL fusion oncoprotein; in the AML cell line MV-4-11, in which Ras is activated downstream of a FLT3 internal tandem duplication; or in the T-cell acute lymphoblastic leukemia cell line CCRF-CEM, which harbors a *KRAS^G12D^* mutation (**Fig. S2D**).

### BET and MEK Inhibition Converge to Co-Target Myc Protein Expression and Myc-Regulated Transcriptional Programs

Whereas BET inhibitors potently suppress Myc transcriptional activity by inhibiting Brd4 ^20,30^, elevated Ras/MAPK signal output results in Myc phosphorylation on serine 62, which stabilizes Myc and promotes the transcription of target genes ^31^. Accordingly, we asked if the synergistic effects of PLX51107 and PD901 in AML cells might be explained by convergent inhibition of Myc protein expression and Myc-regulated transcriptional programs. Exposing OCI-AML3, HL-60, and THP-1 cells to PLX51107 and PD901 resulted in a dose dependent reduction in Myc protein levels (**Fig. 3A**) that were further suppressed by co-treatment (**Fig. 3B**). Furthermore, bulk RNA-seq analysis of *NRAS*-mutant OCI-AML3 cells showed that combining PLX51107 and PD901 strongly potentiated the down-regulation of *MYC* and its transcriptional targets versus either compound alone (**Figs. 3C, S3A, and S3B**) and induced a more immature myeloid state (**Fig. S3C**).

**Figure 3.**
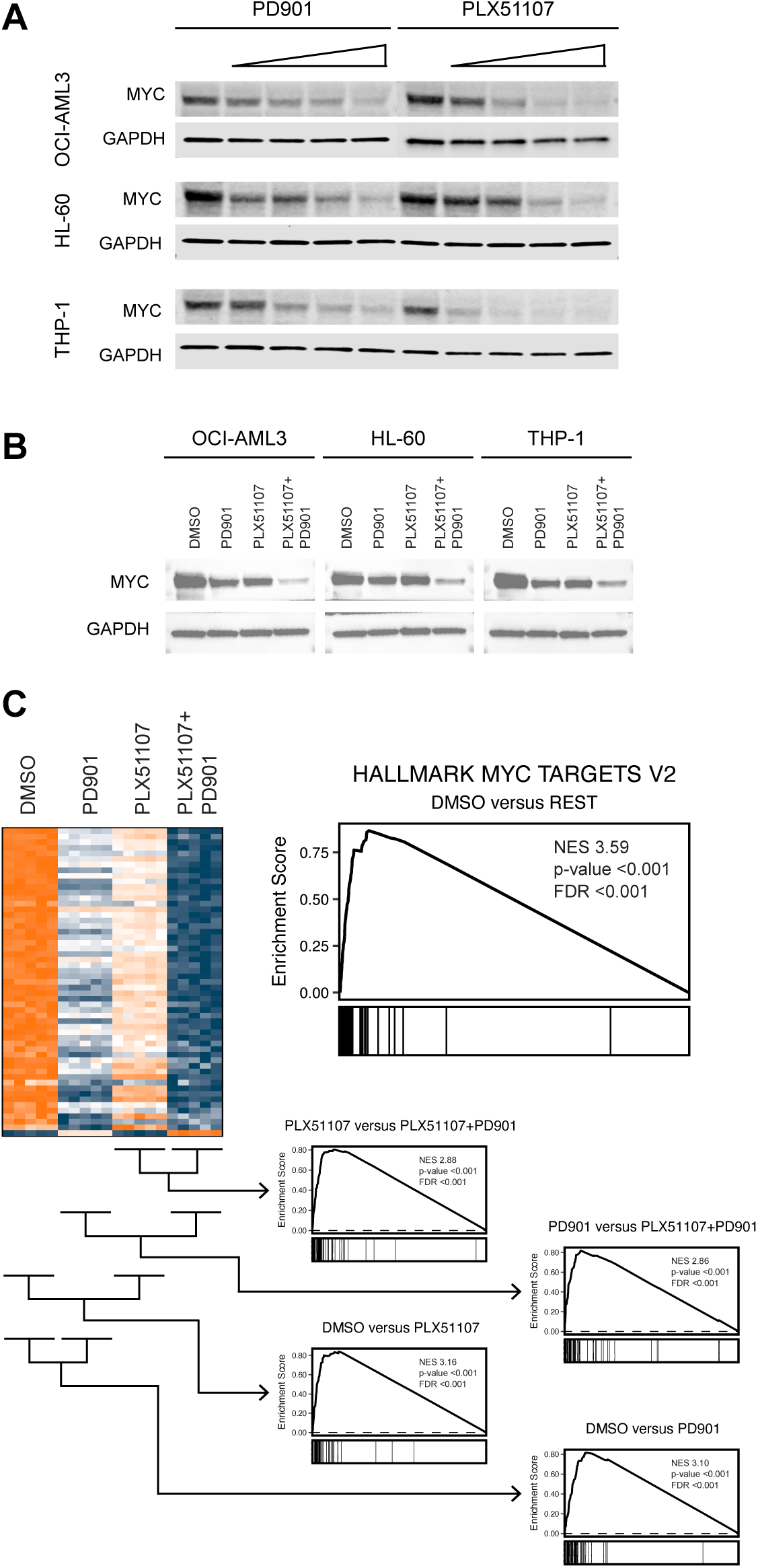
PLX51107 and PD901 induce synergistic down-regulation of Myc protein expression and Myc regulated genes. **A.** Western blotting of lysates prepared from OCI-AML3 (top), HL-60 (middle), and THP-1 (bottom) cells treated with increasing doses of PD901 or PLX51107. **B.** Combination treatment with PLX51107 250 nM and PD901 50 nM potentiates down-regulation of Myc protein levels. **C.** Bulk RNA sequencing (RNA-seq) analysis of OCI-AML3 cells treated with DMSO, PD901 50 nM, PLX51107 250 nM, or the combination reveals suppression of Myc regulated genes with PD901 or PLX51107 that is potentiated with the combination. See also **Figure S3**.

### Efficacy of PLX51107 + PD901 in Primary Murine *Nras^G12D^* AMLs

We previously performed retroviral insertional mutagenesis in mice to generate genetically diverse panels of AMLs characterized by *Nf1* inactivation or endogenous expression of *Nras^G12D^*or *Kras^G12D^* ^32–34^. Transplanting these primary leukemias into mice and treating them *in vivo* is an unbiased strategy for evaluating the efficacy of individual agents or drug combinations and for characterizing mechanisms of response and resistance ^12,33,35–37^. We transplanted mouse *Nras^G12D^* AMLs 6606, 6613, 6695, and 6768 ^12,33,35–37^ into cohorts of recipient mice and treated them with control vehicle, PD901, PLX51107, or PLX51107 + PD901 (n = 5 for each treatment arm). These leukemias have a myelomonocytic/monocytic (M4/M5-like) morphology ^33^ and harbor retroviral integrations in known leukemia oncogenes including *Sox4* (AMLs 6606, 6613, and 6695), *Evi1* (AML 6613), and *Myb* (AML 6768) (**Table S2**). All are biologically aggressive and cause leukocytosis, severe anemia, and death within 2-4 weeks after transplantation into sublethally irradiated congenic recipient mice ^12,33^.

Treatment with PLX51107 (10 mg/kg/day) resulted in 4-fold longer survival compared to vehicle control in recipient mice transplanted with primary *Nras*-mutant AMLs (p-value < 0.0001 compared to vehicle-treated mice) that was further enhanced to 5-fold by the addition of a modest dose of PD901 (1.5 mg/kg given 4 days per week; p-value < 0.0001 compared to vehicle-treated mice and p-value < 0.0083 compared to recipients treated with PLX51107 alone) (**Fig. 4A**). Leukemias 6606, 6695, and 6613 were highly responsive to PLX51107 and/or combination treatment, while AML 6768 was somewhat less sensitive (**Fig. S4A**). As Myb is a key effector of Myc in myeloid cells, constitutive activation due to the *Myb* integration in AML 6768 may underlie this attenuated response to PLX51107. *Ex vivo* studies of primary leukemia cells were consistent with the survival data with the PLX51107 + PD901 combination potently inducing apoptosis, which was most pronounced in the three highly responsive AMLs (**Fig. 4B**).

**Figure 4.**
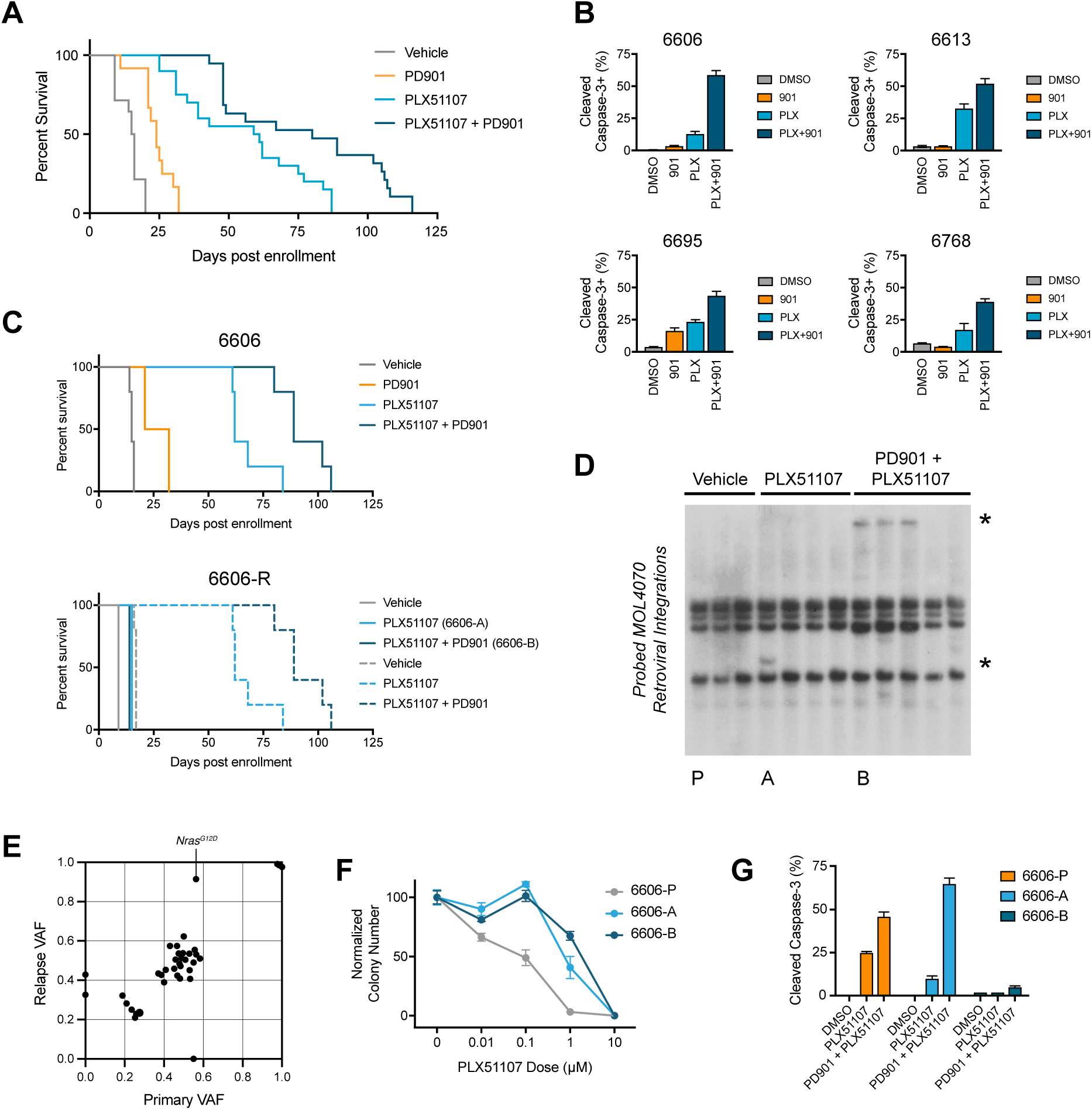
Response and resistance of *Nras^G12D^* AMLs to PLX51107 and PD901. **A.** Kaplan-Meier analysis demonstrates 4-fold median survival extension of recipients transplanted with *Nras^G12D^* AMLs assigned to receive PLX51107 compared to vehicle controls (p < 0.0001 by log-rank test). The addition of low dose PD901 (1.5 mg/kg/day for four days per week) further extended median survival to 5-fold (p < 0.0001 by log-rank test). **B.** *Ex vivo* exposure to both PLX51107 100 nM and PD901 100 nM increases apoptosis in primary AMLs 6606, 6613, 6695, and 6768. **C.** <u>Top</u>. Survival of mice transplanted with AML 6606 treated with vehicle, PD901, PLX51107, or PLX51107 + PD901 (n = 5 per group) (p < 0.0001). <u>Bottom</u>. Relapsed AMLs 6606A and 6606B were harvested from moribund mice, re-transplanted, and re-treated with vehicle or the drug regimen used in the primary preclinical trial (PLX51107 or PLX51107 + PD901). The *in vivo* sensitivity of AMLs 6606A and 6606B is significantly reduced relative to AML 6606 (p < 0.0001 by log-rank test). **D.** Southern blot analysis of DNA isolated from independent recipient mice transplanted with AML 6606 that were treated with either vehicle, PLX51107, or PLX51107 + PD901. Asterisks indicated novel restriction fragments detected with a probe to the MOL4070LTR virus in mice treated with PLX51107 (6606A; lane 4) or PLX51107+PD901 (6606B; lanes 8-10). Note that the same novel restriction fragment is present in relapsed leukemias isolated from the three independent recipients in lanes 8-10, which indicates it emerged from a pre-existing clone. The relapsed leukemias shown in lanes 1 (6606P), 4 (6606A), and 8 (6606B) were expanded *in vivo* and DNA/RNA were isolated for whole exome sequencing (WES) and RNA-seq, respectively. **E.** WES of AMLs 6606P, 6606A, and 6606B revealed an increased in the *Nras^G12D^* mutant allele frequency to ∼100% but did not uncover any candidate resistance mutations. **F.** Normalized colony number formation of bone marrow cells isolated from secondary recipients of AMLs 6606P, 6606A, and 6606B that were exposed to increasing doses of PLX51107 *ex vivo*. **G.** The primary leukemia cells shown in panel F were also exposed to DMSO, PLX51107, or PLX51107 + PD901 and the percentage of cleaved caspase-3 (CC3) cells was assessed by flow cytometry. Note that AML 6606A, which relapsed after single agent treatment with PLX51107 (panel D), is sensitive to combination treatment. By contrast, AML 6606B emerged in recipient mice that received combination treatment and failed to induce CC3 in response to either PLX51107 or PLX51107 + PD901. See also **Figure S4**.

### Myc Protein Expression, Clonal Evolution, and Drug Resistance in Relapsed AMLs

We isolated leukemia cells at euthanasia from mice that relapsed after initially responding to treatment with either PLX51107 or PLX51107 + PD901. Re-transplanting AMLs that relapsed during continuous *in vivo* treatment and re-treating them confirmed intrinsic drug resistance (**Figs. 4C and S4B**) that was frequently characterized by clonal evolution as demonstrated by Southern blot analysis with a probe to the MOL4070 virus (**Fig. 4D and S4C**). Vehicle-treated parental and relapsed leukemias shared multiple common retroviral integrations, which is consistent with the existence of a “founder” clone as observed in diagnosis/relapse human AML pairs ^38,39^. Some relapsed leukemias exhibited MOL4070 insertions that were not seen in the corresponding parental AML (**Figs. 4D and S4C; Table S2**). Whole exome sequencing performed on genomic DNA isolated from parental and resistant AML 6606 revealed an increase in *Nras^G12D^*allele frequency with loss of the wild-type allele due to somatic uniparental disomy ^33^, but no somatic mutations in genes implicated in BET or Ras/MAPK-regulated pathways (**Fig. 4E**). We also verified intrinsic drug resistance *ex vivo* in methylcellulose cultures (**Fig. 4F**) and CC3 assays (**Fig. 4G**). Notably, none of these leukemias previously exhibited clonal evolution or developed intrinsic drug resistance when treated with PD901 at a substantially higher dose (5 mg/kg/day) ^12^.

Studies of HSPCs transduced with oncogenes implicated restoration of Myc protein expression as a key mechanism of adaptive resistance to BET inhibitor treatment ^30,40^. Accordingly, we hypothesized that relapsed *Nras^G12D^* AMLs would exhibit higher Myc protein levels than the corresponding parental leukemia and/or would retain Myc expression upon exposure to PLX51107. Instead, we unexpectedly identified multiple independent relapsed clones that had lower or similar Myc protein levels as the parental leukemia (**Figs. 5A and S5A**). Relapsed AMLs with loss or down-regulation of Myc protein expression had stable *Myc* mRNA levels (**Fig. 5B**) and a lack of compensatory up-regulation of *Mycn* expression (**Fig. S5B**).

**Figure 5.**
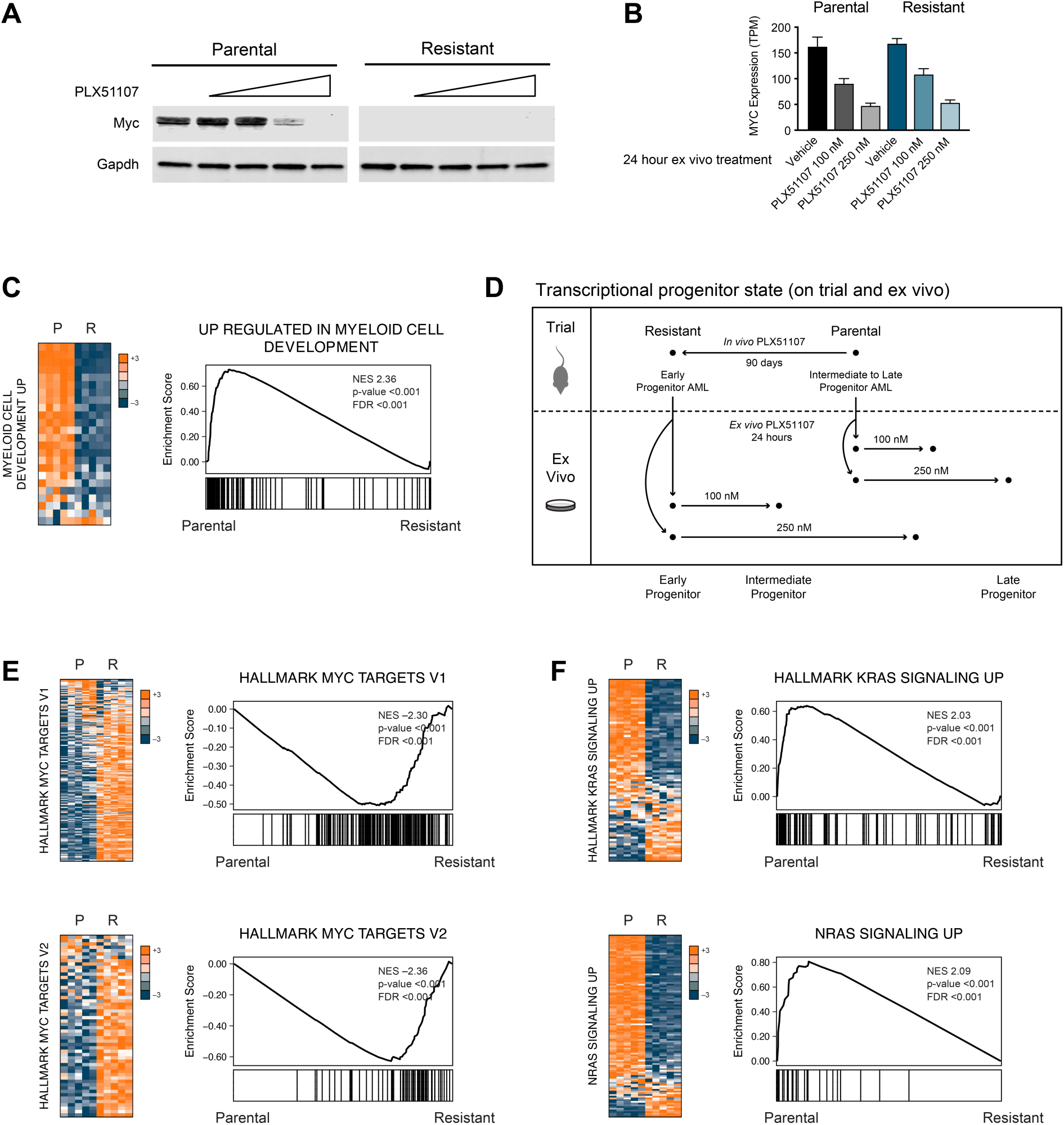
Resistance to PLX51107 is associated with a more primitive transcriptional state and an inverse relationship between Myc- and Ras-associated gene sets. **A.** Western blot analysis of lysates shows the expected down-regulation of Myc protein expression in response to PLX51107 treatment in parental AML 6606P. By contrast, resistant AML 6606B had no detectable Myc protein. B. *Myc* transcript levels are similar in AMLs 6606P and 6606B and decrease in response to increasing doses of PLX51107 *ex vivo*. **C-F.** Bulk RNA-seq analysis of AMLs 6606P and 6606B revealed loss of a myeloid differentiation transcriptional signature, up-regulation of Myc target genes, and down-regulation of oncogene *KRAS* and *NRAS* gene sets in resistant AML 6606B relative to the 6606P. A similar pattern of de-differentiation relative to the corresponding parental AML was also seen in resistant AMLs 6606A, 6613A, and 6613B. See also **Figure S5**.

We performed additional molecular and functional analysis of pairs of relapsed leukemias isolated from recipients of AMLs 6606 and 6613. In these studies, we analyzed control cells isolated from a vehicle-treated recipient transplanted with the respective parental (P) leukemia and from two independent recipients of AMLs 6606 and 6613 that relapsed after drug treatment (A and B). To compare basal and dynamic transcriptional profiles of parental leukemias isolated from vehicle-treated mice (AMLs 6606P and 6613P) and resistant leukemias that emerged after drug treatment (AMLs 6606A/B or AMLs 6613A/B), we performed RNA-seq on parental and resistant counterparts. We also treated parental AML 6606P and resistant leukemia 6606B *ex vivo* with PLX51107 for 24 hours and performed bulk RNA-seq after short-term drug treatment. Consistent with our observations in OCI-AML3 cells (**Fig. S3C**), exposing parental *Nras^G12D^* AML 6606P to PLX51107 resulted in dose-dependent transcriptional changes consistent with myeloid maturation (**Fig. 5D and S5C**). By contrast, near isogenic resistant AML 6606B that emerged at relapse after *in vivo* treatment uniformly expressed relatively more primitive hematopoietic stem cell (HSC)-like transcriptional signatures (**Figs. 5C and S5D**) characterized by up-regulation of Myc transcriptional targets, down-regulation of Ras-associated transcriptional programs (**Figs. 5E-F, S5E-F**), and no enrichment in Wnt/β-catenin signatures (**Fig. S5G**). These observations in AML 6606B were corroborated by analyzing RNA-seq from other relapsed AMLs (**Fig. S5H**). Intriguingly, we found that a similar Myc/Ras pathway relationship occurs during normal hematopoiesis, with normal HSCs preferentially expressing Myc-target genes and more mature myeloid cells preferentially expressing Ras related transcriptional signatures (**Fig. S6**) ^41^. Together with data presented in Figure 4, these results indicate that treatment with PLX51107 selects for the outgrowth of AML clones that are arrested at an immature stage of differentiation and rewire Myc/Ras transcriptional programs independent of Myc protein levels.

### Relapsed Pediatric AMLs Transition to an Earlier Progenitor and MYC Up-regulated State

Whereas clonal evolution has been extensively characterized at the genomic level in adult and pediatric AML, global transcriptional changes are less well studied. Accordingly, we analyzed paired diagnosis-relapse bulk RNA-seq data generated by the NCI TARGET initiative from pediatric AML bone marrow aspirates with tumor purity of greater than 50% at both diagnosis and relapse time points as assessed by flow cytometry (**Fig. 6A**). Intriguingly, *MYC* gene expression did not correlate with the degree of Myc target up-regulation at diagnosis (R^2^ = 0.1490) (**Fig. 6B**), and *MYC* gene expression fold change from diagnosis to relapse did not correlate with changes in MYC target gene expression (R^2^ = 0.1502) (**Fig. 6C**). These findings are consistent with our observations in murine *Nras*-mutant AMLs that were treated with PLX51107 and PD901 (**Figs. 5B and 5E**). Gene expression log ratios were calculated in diagnosis-relapse pediatric AML pairs using HSC, MYC, and Ras gene signatures. This analysis revealed that nearly every leukemia transitioned to a more stem-like state at relapse that was associated with elevated expression of MYC target genes (**Figs. 6D and 6E**) without concomitant changes in Wnt/β-catenin signatures (**Fig. S7**). A subset of cases characterized by monocytic differentiation at diagnosis (FAB M2-M5) also showed decreased expression of gene sets associated with oncogenic Ras signaling at relapse (**Fig. 6F**).

**Figure 6.**
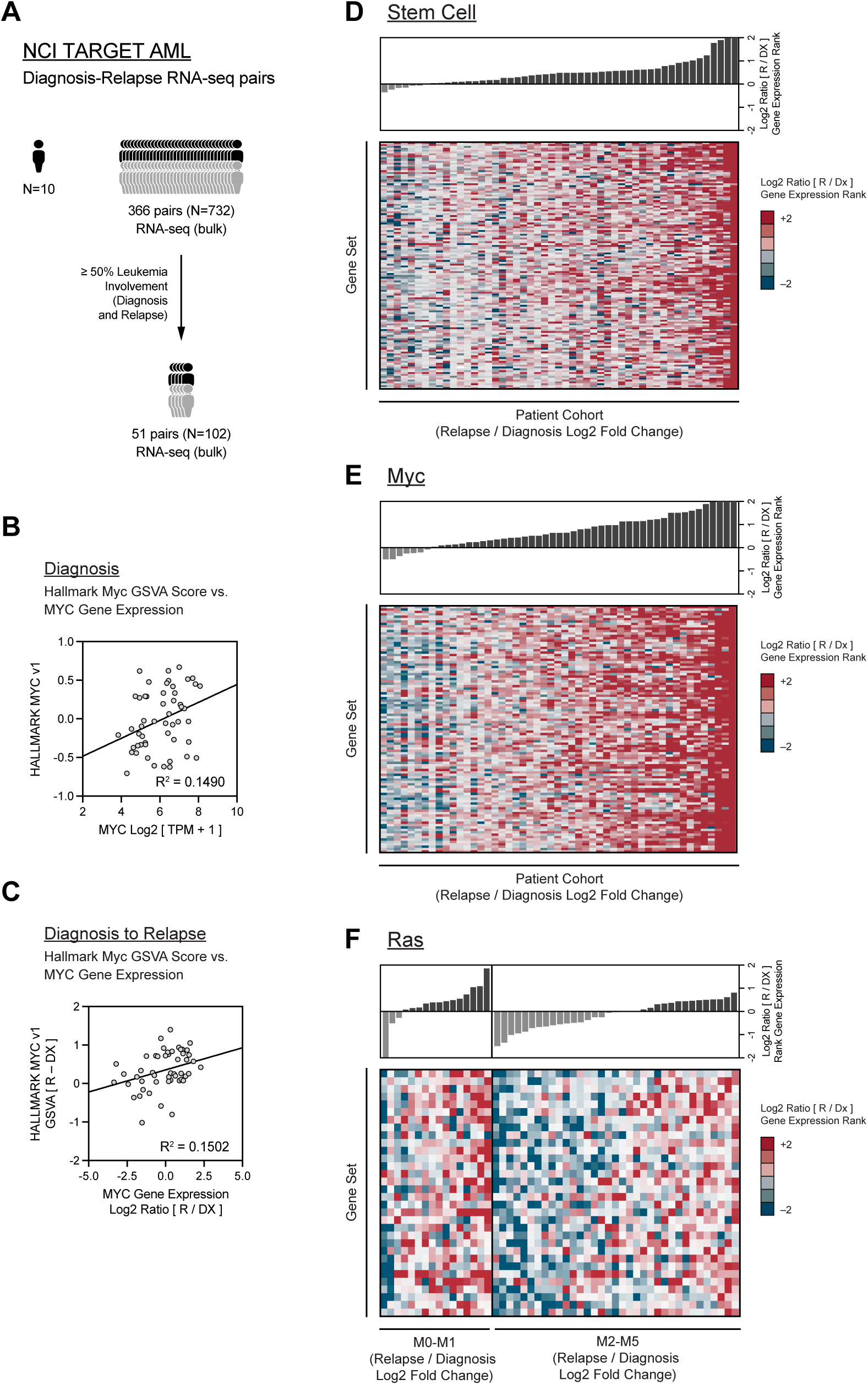
Relapsed pediatric AMLs are enriched for immature and MYC associated transcriptional programs. **A.** NCI TARGET AML transcriptome data were analyzed, with a specific focus on diagnosis-relapse pairs (N = 732 or 366 pairs). Bulk RNA sequencing (RNA-seq) analysis was restricted to diagnosis-relapse pairs with 50% or more leukemia involvement at both time points (N = 102 or 51 pairs) to optimize signal-to-noise ratio between normal bone marrow and leukemia cell populations. **B, C** Neither *MYC* gene expression at diagnosis nor fold change from diagnosis to relapse are correlated with MYC target gene expression. **D-E.** Immature morphologic AML subtypes (FAB M0 and M1) at diagnosis exhibited up-regulation of MYC and stem-like transcriptional signatures at relapse. More mature morphologic AML subtypes (FAB M2-M5) were associated with the same stem/MYC transitions and also down-regulation of Ras transcriptional signatures. See also **Figure S7**.

We next analyzed single cell RNA and ATAC sequencing (scRNA/ATAC-seq) data from a cohort of pediatric AML cases profiled at diagnosis and relapse ^42^. Based on our recent finding that *KMT2A*-rearranged AMLs present with monocytic transcriptional programs at diagnosis ^42^, we focused on this this subtype (N = 7). Interrogating the Broad Molecular Signature Database revealed similar transcriptional transitions at the single cell level (**Figs. 7A, 7B, and S8**).

**Figure 7.**
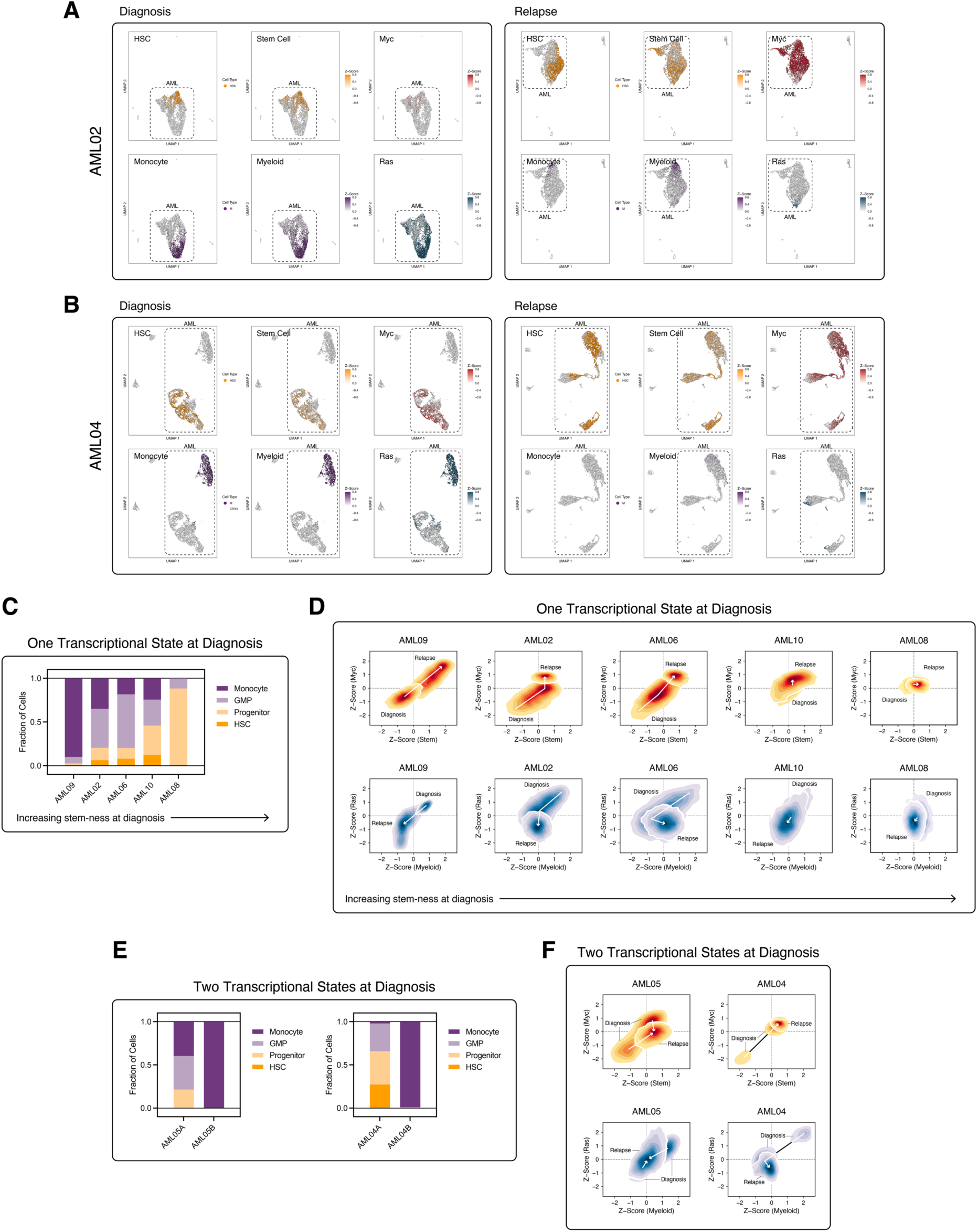
Single cell RNA and ATAC sequencing show analogous patterns of transcriptional plasticity and provide insights into intra-tumoral transcriptional heterogeneity. A, **B.** Single cell RNA and ATAC sequencing (scRNA/ATAC-seq) of diagnosis and relapsed KMT2A-rearranged AML02 and AML04 reveals transitions towards upregulation of MYC and stem-like transcriptional programs and down-regulation of Ras and myeloid transcriptional programs. HSC and Monocyte UMAP plots were assigned a color based on the assigned hematopoiesis maturation as described in Lambo, et al.^42^ Stem Cell, Myeloid, Myc, and Ras UMAP plots were shaded based on gene signature Z-score expression from Broad Molecular Signature Database gene sets. **C, D.** Five AMLs are associated with one transcriptional state at diagnosis (similar to AML02). Two-dimensional contour plot displaying the relative transcriptional transitions comparing MYC versus Stem and Ras versus Myeloid signatures. **E, F.** The other two AMLs are associated with two distinct transcriptional states at diagnosis (similar to AML04), one that is more primitive and one that is more mature. See also **Figure S8**.

Specifically, five of seven AMLs uniformly transitioned towards a more primitive, MYC target up-regulated state with concomitant down-regulation of Ras transcriptional programs. Furthermore, the degree of transition was proportional to the fraction of cells that were HSC-like at diagnosis (**Figs. 7C and 7D**). The other two leukemias (AML04 and AML05) each had two transcriptionally distinct cell populations present at diagnosis (**Figs. 7E and 7F).** These two transcriptional states coalesced towards the more primitive, MYC up-regulated state at relapse, with relapse occurring more rapidly than in the other five AMLs.

## DISCUSSION

High throughput sequencing studies uncovered pronounced differences in the mutational profiles of AMLs arising in pediatric, young adult, and elderly individuals ^1,3^. In older adults, AML is primarily initiated by somatic mutations in *DNMT3A, TET2, ASXL1,* and other “epigenetic” regulators ^43^ with secondary mutations in genes encoding key signaling molecules such as *FLT3* and *NRAS* cooperating to drive leukemic transformation ^44^. *TP53* mutations occur in ∼10% of adult AMLs and portend particularly poor outcomes. By contrast, pediatric AML is characterized by a lower overall mutational burden, fewer epigenetic and *TP53* mutations, more frequent gene fusions of *KTM2A* and other transcription factors, and a high prevalence of *FLT3, NRAS,* and *KRAS* mutations ^1^. The low incidence of *DNMT3A, TET2,* and *ASXL1* mutations in pediatric AML suggests that the underlying epigenetic landscapes differ substantially in pediatric and adult AMLs. This, in turn, may strongly modulate the responses to PLX51107 and other chemical inhibitors that broadly target transcriptional networks regulating cell identity, differentiation, and survival.

Consistent with the elegant *in vivo* shRNA screen that identified Brd4 as a dependency in mouse cells engineered to express *KTM2A-AF9* and *NRAS^G12D^* ^20^, we show that PLX51107 is efficacious in primary mouse leukemias with similar molecular and biologic features to pediatric AML ^33^. We further show that BET and MEK inhibition synergistically suppress AML growth and Myc transcriptional output. These data are consistent with a recent analysis of the BEAT AML dataset that associated a transcriptional profile corresponding to monocytic differentiation with *in vitro* sensitivity to MEK and BET inhibitors ^25^. Our preclinical studies co-targeting BET bromodomain proteins and aberrant signal transduction in a mouse model of hematologic cancer are reminiscent of a similar approach in the *Mpl^V515L^* mouse model of myelofibrosis that was successfully translated clinically ^24^.

We provide new mechanistic insights that extend the known relationship of the anti-leukemia activity of BET inhibition with Myc suppression ^20,30^. Specifically, we find that resistant AMLs that emerge after continuous *in vivo* treatment up-regulate both Myc and stem-like transcriptional programs irrespective of Myc protein levels and simultaneously down-regulate Ras-associated transcriptional programs. These data are unexpected as short-term PLX51107 exposure promoted myeloid differentiation in AML cell lines *in vitro* and primary mouse AMLs *ex vivo*. Two groups transduced mouse HSPCs with *KMT2A::AF9* to characterize mechanisms of adaptative resistance to BET inhibition *in vitro* ^30,40^. In agreement with our results, these studies uncovered sustained Myc gene expression and, in one study, transition to a more stem cell-like phenotype in resistant cells. Whereas both groups implicated Wnt/β-catenin activation in resistance, our integrated analysis support a dominant role of increased Myc target transcriptional output that is independent of Wnt/β-catenin up-regulation in primary *Nras*-mutant mouse AMLs and in pediatric leukemias that relapsed after BET inhibition and conventional front-line chemotherapy, respectively ^42^. One potential explanation for these differences is that we investigated primary AMLs with diverse initiating mutations that were treated *in vivo* while these previous studies utilized mouse HSPCs transduced with *KMT2A::AF9* vectors and resistance that emerged *in vitro* ^30,40^.

The pattern of response and resistance to PLX51107 has interesting parallels with recent studies of venetoclax, which is a front-line therapy for older AML patients who are unable to tolerate intensive chemotherapy ^45^. Pei, et al. recently reported that treatment with venetoclax and azacitidine drives AML to a more differentiated state that is characterized by the emergence of drug resistant clones harboring *NRAS* mutations ^26,27^. Resistance to PLX51107 in *Nras*-mutant AMLs with monocytic features is similarly associated with global transcriptional changes; however, these occur in the opposite direction across the myeloid differentiation spectrum with PLX51107 resistant AMLs adopting more HSC-like identities. Transcriptional plasticity in AML has potential therapeutic implications and may explain, in part, the disappointing efficacy of CAR-T cell-based therapies to date as many target antigens are expressed at discrete developmental stages. The transcriptional transitions across the myeloid differentiation spectrum observed in response to venetoclax ^26,27^ and PLX51107 also raises the intriguing possibility that combining these drugs might enhance survival by suppressing a key mechanism of resistance to each single agent. In a recent trial of PLX51107 and azacitidine in relapsed and refractory AML, 7 of 8 patients with an objective clinical response had progressed after prior venetoclax treatment ^22^.

An advantage of the transplantable primary AML models utilized in this study is the ability to compare data from serial preclinical trials of single agents and drug combinations. We previously treated the four *Nras*-mutant leukemias included in the present study with PD901 at a daily dose of 5 mg/kg/day (35 mg/kg/week). Consistent with studies of allosteric MEK inhibitors in human AML ^9^, PD901 treatment caused cell cycle arrest and resulted in a modest increase in survival without clonal evolution or adaptive resistance ^12^. By contrast, a much lower, intermittent dose of PD901 (6 mg/kg/week) exhibited significant synergistic activity with PLX51107 in down-regulating Myc protein levels and transcriptional output and significantly extended survival in comparison to PLX51107 alone. We speculate that MEK inhibition blocks a key survival pathway that AML cells deploy in response to BET bromodomain inhibition.

Taken together, our mechanistic and preclinical data provide a compelling rationale for evaluating MEK and BET inhibitor combinations in AML, particularly in leukemias with *NRAS* mutations and/or evidence of monocytic differentiation. PLX51107 and similar BET inhibitors might also enhance the anti-leukemia activity of venetoclax and of the emerging class of menin inhibitors, which have also been shown to down-regulate Myc transcriptional output ^46^.

## METHODS

### Cell line authentication and quality control

Cell lines were obtained from the American Type Culture Collection (ATCC) or Deutsche Sammlung von Mikroorganismen und Zellkulturen (DSMZ), expanded, and stored at early passage. Early passage OCI-AML3, HL-60, and THP-1 cells (DSMZ and ATCC) were cultured in RPMI media (HyClone) containing 10% heat inactivated fetal bovine serum (Corning), 1% penicillin-streptomycin (Thermo Fisher), and 1% GlutaMAX^TM^ (Thermo Fisher) at 37°C and in 5% CO2. During passage, the cells were tested for mycoplasma regularly with a mycoplasma PCR detection kit (Millipore Sigma). STR profiling was performed once and compared with external STR profiles of cell lines (when available) to determine cell line ancestry.

### *Nras^G12D^* murine acute myeloid leukemias

F1 C57BL/6 × 129Sv/Jae strain background ^12,33^ was utilized for the described acute myeloid leukemia (AML) animal experiments. Each AML was expanded *in vivo*, and the same number of cells (2 x 10^6^) was injected into recipient mice, which were randomly assigned to treatment or control arms. Importantly, the time to death for vehicle-treated mice transplanted with individual leukemias was highly consistent across serial studies. The identities of individual AMLs and relationship to resistant clones were verified by molecular fingerprinting (Southern blot). Data analysis from previous studies has verified the statistical power of the cohort sizes used in these trials to reliably detect significant differences ^12,32,37,47^. Littermates were group housed, provided free access to standard rodent diet and water, and were randomly assigned to experimental treatment groups. For therapeutic studies, primary cryopreserved AML cells (2 x 10^6^) were injected intravenously by tail vein into 8-12 week old mice that received a sublethal radiation dose (600 rads). Primary bone marrow from moribund mice was then serially passaged using the same methods into a cohort of recipient animals for assignment to therapeutic cohorts. Animals randomly assigned to experimental treatment groups were dosed by oral gavage without blinding at any stage of the study. All animals assigned to treatment groups were included in the survival analysis. Welfare-related assessments and interventions were carried our daily during the treatment period. All studies were approved by the Institutional Animal Care and Use Committee at the University of California, San Francisco.

### Primary AML cell culture

For short-term cultures of primary murine AML cells, bone marrow cells from moribund transplant recipients were plated at 5×104 cells per mL in AML medium [IMDM supplemented with 10% fetal bovine serum, penicillin/streptomycin, glutamine, SCF (10 ng/mL), GM-CSF (10 ng/mL), and IL-3 (8 ng/mL)] and grown at 37°C. Cultured cells were grown with three or more technical replicates for each condition.

### NCI TARGET

AML samples were collected with informed consent from patients (0-28 years of age) enrolled on Children’s Oncology Group (COG) trials AAML0531 (NCT00372593), and AAML1031 (NCT01371981). IRB approval for each protocol was obtained at each participating institution and submitted to the CTSU regulatory office.

### Cell viability analysis

Cell lines were plated at 1,000 cells/well on white opaque 96-well flat bottom plates (Perkin Elmer) in technical triplicate. Cells were treated with indicated drug dose series for 72 hours. Metabolically active cells as a proxy for overall cell viability were measured with CellTiter-Glo (Promega) and analyzed on a plate reader (Tecan M2000 Infinite Pro).

Experiments were performed with technical triplicates and a representative experiment is shown from at least three experimental replicates for each cell line.

### Apoptosis analysis

Cell lines were plated at 20,000 cells/well on 96-well round bottom plates (Corning) in triplicates. Cells were treated with indicated drug dose series for 48 hours. Cells were collected at the bottom of the plates and fixed with 4% paraformaldehyde (Electron Microscopy Sciences) at room temperature in the dark for 10 minutes. After collection, cells were permeabilized with ice cold methanol for 30 minutes on ice. Cells were then washed with ice cold PBS (Thermo Fisher) twice, followed by one wash with FACS buffer (PBS containing 2% fetal bovine serum). Cells were stained with cleaved caspase-3 antibody (BD Horizon 560627) at 1:40 dilution for one hour, and subsequently washed with FACS buffer three times before analyzing on an Attune NxT flow cytometer (Thermo Fisher). Experiments were performed with technical triplicates and a representative experiment is shown from at least three experimental replicates for each cell line.

### MOL4070LTR Integration Cloning

Restriction enzyme digestion of genomic DNA from mouse AMLs, gel electrophoresis, Southern blot analysis, hybridization with a MOL4070LTR-specific probe was performed as described previously ^12,32,36^. Junctional fragments at sites of retroviral integration were identified as previously described using linker-based PCR amplification and sequencing ^12,32,36^.

### Immunoblotting

Cells were lysed in ice cold RIPA buffer (Pierce) supplemented with HaltTM protease and phosphatase inhibitor cocktail (Thermo Fisher) and 0.5µM EDTA, incubated on ice for 30 minutes, and centrifuged at maximum speed for 15 minutes to collect whole cell lysates. Protein concentration was measured with the BCA protein assay (Pierce). 30µg of total protein per sample was loaded into 4-12% gradient Criterion^TM^ TGXTM gels (Bio-Rad) and separated by SDS-PAGE. Proteins were transferred to PVDF membranes. c-Myc antibody (Cell Signaling Technology Cat# 5605, RRID:AB_1903938) was used for detecting c-Myc and Hsp90 antibody (BD Biosciences Cat# 610418, RRID:AB_397798) was used for loading controls.

### Real time quantitative real-time polymerase chain reaction

Total RNA was extracted from cell line or primary bone marrow cells using RNeasy RNA Extraction Kits (Qiagen). Reverse transcription was performed using SuperScript Reverse Transcriptase (Invitrogen). RT-qPCR were setup using TaqMan Gene Expression Assays (ThermoFisher) and the indicated gene-specific probe and primer sets and run on a QuantStudio 5 (ThermoFisher). Experiments were performed with technical triplicates and a representative experiment is shown from three experimental replicates for each probe.

### Whole exome sequencing

Genomic DNA was extracted from the bone marrow cells and then sheared to generate 150 to 200 base pair fragments using a Covaris S2 focused-ultrasonicator. Indexed libraries were prepared using the Agilent SureSelect XT2 Reagent Kit for the HiSeq platform. Exomes were captured using the Agilent SureSelect XT2 Mouse All Exon bait library. Sample quality and quantity were assessed using the Agilent 2100 Bioanalyzer instrument.

Paired-end 100 base pair reads were generated on an Illumina HiSeq 2000 platform. All sequence data including read alignment; quality and performance metrics; post-processing, somatic mutation and DNA copy number alteration detection; and variant annotation were performed as previously described ^48^ using the mm10 build of the mouse genome. Briefly, reads were aligned with Burrows-Wheeler Aligner, and processed using Picard (http://broadinstitute.github.io/picard) tools and the Genome Analysis Toolkit (GATK) to perform base quality recalibration and multiple sequence realignment. Single nucleotide variants and indels were detected with the MuTect and Pindel algorithms, respectively. Candidate somatic mutations were manually reviewed using Integrative Genomics Viewer.

### Bulk RNA library construction, sequencing, and analysis

Total RNA extracted from peripheral blood or bone marrow diagnostic specimens was purified using AllPrep DNA/RNA/miRNA Universal Kits (Qiagen). Purified RNA samples were then prepared for either strand-specific polyadenylated enriched messenger RNA libraries or strand-specific ribosome RNA-depleted libraries by the British Columbia Genome Sciences Center (BCGSC). Paired-end sequencing was performed on Illumina HiSeq 2000/2500 platforms and sequence reads were aligned to the GRCh37 reference genome using BWA (v0.5.7). Reads were discarded based on mapping quality or if they failed the Illumina chastity filter and duplicate reads were marked using Picard (v1.11). Gene level coverage analysis was performed using the BCGSC pipeline v1.1 with Ensembl v69 annotations and was normalized based on RPKM or TPM. Gene Set Enrichment Analysis (GSEA) and Gene Set Variation Analysis (GSVA) was performed using the Broad Molecular Signature Database (across the entire Hallmark, C2, and C6 collections).

Notable gene sets that were frequently significant in our analysis included Hallmark Myc Targets v1 and v2, Hallmark Kras Signaling Up, Croonquist Nras Signaling Up, Jaatinen Hematopoietic Stem Cell Up, the Ivanova series (e.g., Ivanova Hematopoiesis Stem Cell), Brown Myeloid Cell Development Up and Down, among many others highlighted in **Figures S5 and S6**.

### Single cell RNA and ATAC library construction, sequencing, and data processing

Processing, sequencing, and analysis was performed as previously described ^42^. Briefly, for scRNA-seq, single-cell 3′ RNA-seq libraries were generated using the Chromium Single Cell 3′ Library & Gel Bead Kit v3.0 (10X Genomics) following the manufacturer’s protocol. Prior to nuclei isolation for scATAC-seq, CD15 magnetic bead-based depletion was performed using a combination of magnetic streptavidin beads and a biotin-conjugated CD15 monoclonal antibody, to increase data quality. This protocol depletes neutrophils, which have low quality ATAC profiles, while retaining important cellular hierarchies within peripheral blood mononuclear cells. Based on pilot experiments comparing CD15 depleted and undepleted data from the same sample, no reduction of either malignant cells or inferred myeloid cell populations was detected^42^. Nuclei isolation was subsequently performed according to instructions in the 10x Genomics Demonstrated Protocol CG000169. Single-cell ATAC-seq libraries were then generated using the Chromium Single Cell ATAC Reagent Kit v1 or Chromium NextGEM Single Cell ATAC Reagent Kit v1.1 (10X Genomics). scRNA and scATAC library quality was assessed using the Agilent High Sensitivity DNA kit and libraries were quantified using the Qubit dsDNA HS Assay kit (Thermo Fisher Scientific). Libraries were pooled and converted using the MGIEasy Universal Library Conversion Kit (App-A; MGI Americas) following the manufacturer’s protocol and as described above. Converted libraries were sequenced using the DNBSEQ-G400RS high-throughput sequencing set on the DNBSEQ-G400 sequencer using 10X sequencing parameters. Processing of scRNA-seq and scATAC-seq data was performed as described in Lambo, et al. ^42^. KMT2A-rearranged AML cases were selected based the following requirements: (1) availability of diagnosis and relapse timepoints (omission of AML01 and AML03); and, (2) adequate number of cells at both diagnosis and relapse timepoints (N = 500 malignant cells) (omission of AML07 and AML11). Applied Broad Molecular Signature Database gene sets are described in the bulk RNA methods section.

### Quantification and Statistical Analysis

Quantification and statistical analysis were performed with either GraphPad Prism (GraphPad) or the open-source R Statistical Computing software (http://www.r-project.org/) utilizing the statistical tests described in the text and figure legends. In animal experiments, n represents number of animals utilized in each treatment group, and survival analysis is represented as a Kaplan-Meier analysis with statistical significance calculated utilizing the log-rank test.

## Acknowledgements

This work was supported by grants from the National Institutes of Health (K08 CA256489 [B.J.H.], R01 CA193994 [K.S.], and U54 CA196519 [K.S.]), by the Leukemia and Lymphoma Society (Translational Research Program Award 6612-20 [K.S.]), and by a St. Baldrick’s Foundation (Fellowship Award [B.J.H]). This work utilized the Helen Diller Family Comprehensive Cancer Center core facilities in Computational Biology and Informatics and Laboratory for Cell Analysis. The results published here are in part based on data generated under the Therapeutically Applicable Research to Generate Effective Treatments (TARGET) project managed by the National Cancer Institute. Information about TARGET can be found at https://ocg.cancer.gov.

## Authorship Contribution

Conceptualization, B.J.H., K.S.; Methodology, B.J.H., G.B., K.S.; Investigation, B.J.H., J.D., A.B., M.D.H., X.W., E.H., A.M.W., M.R.B.; Formal Analysis, B.J.H.; Resources, M.R.B., Q.L., S.M., G.B., K.S.; Writing, B.J.H., K.S.; Revisions, all authors; Supervision, S.M., G.B., K.S.; Funding Acquisition, B.J.H., S.M., K.S.

## Data Availability Statement

Raw sequencing FASTQ files are available at the DNA Data Bank of Japan (DDBJ: PRJDB17372 for mouse whole exome sequencing data), NCBI Gene Expression Omnibus (GEO: GSE254173 for mouse bulk RNA sequencing data), NCBI Database of Genotypes and Phenotypes (dbGaP: https://www.ncbi.nlm.nih.gov/gap under study ID phs000465.v21.p8 for human bulk RNA sequencing data), and as previously described 58 for human single cell RNA and ATAC sequencing data.

## Notes

**Conflict of Interest Disclosure Statement** Gideon Bollag is the CSO of Opna Bio and was previously CEO of Plexxikon. Michael R. Burgess is an employee of Bristol Myers Squibb. The remaining authors declare no financial interest related to this work.

### Competing Interest Statement

Gideon Bollag is the CSO of Opna Bio and was previously CEO of Plexxikon. Michael R. Burgess is an employee of Bristol Myers Squibb. The remaining authors declare no financial interest related to this work.

## REFERENCES

1 Bolouri, H. et al. The molecular landscape of pediatric acute myeloid leukemia reveals recurrent structural alterations and age-specific mutational interactions. Nat Med 24, 103–112 (2018). 10.1038/nm.4439

2 Lindsley, R. C. & Ebert, B. L. The biology and clinical impact of genetic lesions in myeloid malignancies. Blood 122, 3741–3748 (2013). 10.1182/blood-2013-06-460295

3 Papaemmanuil, E. et al. Genomic Classification and Prognosis in Acute Myeloid Leukemia. N Engl J Med 374, 2209–2221 (2016). 10.1056/NEJMoa1516192

4 McMahon, C. M. et al. Clonal Selection with RAS Pathway Activation Mediates Secondary Clinical Resistance to Selective FLT3 Inhibition in Acute Myeloid Leukemia. Cancer Discov 9, 1050–1063 (2019). 10.1158/2159-8290.CD-18-1453

5 Stein, E. M. et al. Enasidenib in mutant IDH2 relapsed or refractory acute myeloid leukemia. Blood 130, 722–731 (2017). 10.1182/blood-2017-04-779405

6 DiNardo, C. D. et al. Molecular patterns of response and treatment failure after frontline venetoclax combinations in older patients with AML. Blood 135, 791–803 (2020). 10.1182/blood.2019003988

7 Samra, B., Konopleva, M., Isidori, A., Daver, N. & DiNardo, C. Venetoclax-Based Combinations in Acute Myeloid Leukemia: Current Evidence and Future Directions. Front Oncol 10, 562558 (2020). 10.3389/fonc.2020.562558

8 Zhang, Q. et al. Activation of RAS/MAPK pathway confers MCL-1 mediated acquired resistance to BCL-2 inhibitor venetoclax in acute myeloid leukemia. Signal Transduct Target Ther 7, 51 (2022). 10.1038/s41392-021-00870-3

9 Borthakur, G. et al. Activity of the oral mitogen-activated protein kinase kinase inhibitor trametinib in RAS-mutant relapsed or refractory myeloid malignancies. Cancer 122, 1871–1879 (2016). 10.1002/cncr.29986

10. Jain, N., et al. Phase II study of the oral MEK inhibitor selumetinib in advanced acute myelogenous leukemia: a University of Chicago phase II consortium trial. Clin Cancer Res 20, 490–498 (2014). 10.1158/1078-0432.CCR-13-1311

11 Hallin, J. et al. The KRAS(G12C) Inhibitor MRTX849 Provides Insight toward Therapeutic Susceptibility of KRAS-Mutant Cancers in Mouse Models and Patients. Cancer Discov 10, 54–71 (2020). 10.1158/2159-8290.CD-19-1167

12 Burgess, M. R. et al. Preclinical efficacy of MEK inhibition in Nras-mutant AML. Blood 124, 3947–3955 (2014). 10.1182/blood-2014-05-574582

13 Zhao, Y. et al. Diverse alterations associated with resistance to KRAS(G12C) inhibition. Nature 599, 679–683 (2021). 10.1038/s41586-021-04065-2

14 Awad, M. M. et al. Acquired Resistance to KRAS(G12C) Inhibition in Cancer. N Engl J Med 384, 2382–2393 (2021). 10.1056/NEJMoa2105281

15 Wang, T. et al. Gene Essentiality Profiling Reveals Gene Networks and Synthetic Lethal Interactions with Oncogenic Ras. Cell 168, 890–903 e815 (2017). 10.1016/j.cell.2017.01.013

16 Lou, K. et al. KRAS(G12C) inhibition produces a driver-limited state revealing collateral dependencies. Sci Signal 12 (2019). 10.1126/scisignal.aaw9450

17 Trojer, P. Targeting BET Bromodomains in Cancer. Annual Review of Cancer Biology 6, 313–336 (2022). 10.1146/annurev-cancerbio-070120-103531

18 Neff, T. et al. Polycomb repressive complex 2 is required for MLL-AF9 leukemia. Proc Natl Acad Sci U S A 109, 5028–5033 (2012). 10.1073/pnas.1202258109

19 Zuber, J. et al. Mouse models of human AML accurately predict chemotherapy response. Genes Dev 23, 877–889 (2009). 10.1101/gad.1771409

20 Zuber, J. et al. RNAi screen identifies Brd4 as a therapeutic target in acute myeloid leukaemia. Nature 478, 524–528 (2011). 10.1038/nature10334

21 Berthon, C. et al. Bromodomain inhibitor OTX015 in patients with acute leukaemia: a dose-escalation, phase 1 study. Lancet Haematol 3, e186–195 (2016). 10.1016/S2352-3026(15)00247-1

22 Senapati, J. et al. Phase I Results of Bromodomain and Extra-Terminal Inhibitor PLX51107 in Combination with Azacitidine in Patients with Relapsed/Refractory Myeloid Malignancies. Clin Cancer Res 29, 4352–4360 (2023). 10.1158/1078-0432.CCR-23-1429

23 Mascarenhas, J. et al. MANIFEST: Pelabresib in Combination With Ruxolitinib for Janus Kinase Inhibitor Treatment-Naive Myelofibrosis. J Clin Oncol 41, 4993–5004 (2023). 10.1200/JCO.22.01972

24 Kleppe, M. et al. Dual Targeting of Oncogenic Activation and Inflammatory Signaling Increases Therapeutic Efficacy in Myeloproliferative Neoplasms. Cancer Cell 33, 29–43 e27 (2018). 10.1016/j.ccell.2017.11.009

25 Bottomly, D. et al. Integrative analysis of drug response and clinical outcome in acute myeloid leukemia. Cancer Cell 40, 850–864 e859 (2022). 10.1016/j.ccell.2022.07.002

26 Pei, S. et al. A novel type of monocytic leukemia stem cell revealed by the clinical use of venetoclax-based therapy. Cancer Discov (2023). 10.1158/2159-8290.CD-22-1297

27 Pei, S. et al. Monocytic Subclones Confer Resistance to Venetoclax-Based Therapy in Patients with Acute Myeloid Leukemia. Cancer Discov 10, 536–551 (2020). 10.1158/2159-8290.CD-19-0710

28 Bowen, D. T. et al. RAS mutation in acute myeloid leukemia is associated with distinct cytogenetic subgroups but does not influence outcome in patients younger than 60 years. Blood 106, 2113–2119 (2005). 10.1182/blood-2005-03-0867

29 Ozer, H. G. et al. BRD4 Profiling Identifies Critical Chronic Lymphocytic Leukemia Oncogenic Circuits and Reveals Sensitivity to PLX51107, a Novel Structurally Distinct BET Inhibitor. Cancer Discov 8, 458–477 (2018). 10.1158/2159-8290.CD-17-0902

30 Fong, C. Y. et al. BET inhibitor resistance emerges from leukaemia stem cells. Nature 525, 538–542 (2015). 10.1038/nature14888

31 Farrell, A. S. & Sears, R. C. MYC degradation. Cold Spring Harb Perspect Med 4 (2014). 10.1101/cshperspect.a014365

32 Lauchle, J. O. et al. Response and resistance to MEK inhibition in leukaemias initiated by hyperactive Ras. Nature 461, 411–414 (2009). 10.1038/nature08279

33 Li, Q. et al. Hematopoiesis and leukemogenesis in mice expressing oncogenic NrasG12D from the endogenous locus. Blood 117, 2022–2032 (2011). 10.1182/blood-2010-04-280750

34 Dail, M. et al. Mutant Ikzf1, KrasG12D, and Notch1 cooperate in T lineage leukemogenesis and modulate responses to targeted agents. Proc Natl Acad Sci U S A 107, 5106–5111 (2010). 10.1073/pnas.1001064107

35 Fenouille, N. et al. The creatine kinase pathway is a metabolic vulnerability in EVI1-positive acute myeloid leukemia. Nat Med 23, 301–313 (2017). 10.1038/nm.4283

36 Burgess, M. R. et al. KRAS Allelic Imbalance Enhances Fitness and Modulates MAP Kinase Dependence in Cancer. Cell 168, 817–829 e815 (2017). 10.1016/j.cell.2017.01.020

37 Wandler, A. M. et al. Loss of glucocorticoid receptor expression mediates in vivo dexamethasone resistance in T-cell acute lymphoblastic leukemia. Leukemia 34, 2025–2037 (2020). 10.1038/s41375-020-0748-6

38 Peretz, C. A. C. et al. Single-cell DNA sequencing reveals complex mechanisms of resistance to quizartinib. Blood Adv 5, 1437–1441 (2021). 10.1182/bloodadvances.2020003398

39 Ding, L. et al. Clonal evolution in relapsed acute myeloid leukaemia revealed by whole-genome sequencing. Nature 481, 506–510 (2012). 10.1038/nature10738

40 Rathert, P. et al. Transcriptional plasticity promotes primary and acquired resistance to BET inhibition. Nature 525, 543–547 (2015). 10.1038/nature14898

41 Novershtern, N. et al. Densely interconnected transcriptional circuits control cell states in human hematopoiesis. Cell 144, 296–309 (2011). 10.1016/j.cell.2011.01.004

42 Lambo, S. et al. A longitudinal single-cell atlas of treatment response in pediatric AML. Cancer Cell (2023). 10.1016/j.ccell.2023.10.008

43 Xie, M. et al. Age-related mutations associated with clonal hematopoietic expansion and malignancies. Nat Med 20, 1472–1478 (2014). 10.1038/nm.3733

44 Welch, J. S. et al. The origin and evolution of mutations in acute myeloid leukemia. Cell 150, 264–278 (2012). 10.1016/j.cell.2012.06.023

45 DiNardo, C. D. & Wei, A. H. How I treat acute myeloid leukemia in the era of new drugs. Blood 135, 85–96 (2020). 10.1182/blood.2019001239

46 Wu, G. et al. Menin enhances c-Myc-mediated transcription to promote cancer progression. Nat Commun 8, 15278 (2017). 10.1038/ncomms15278

47 Dail, M. et al. Loss of oncogenic Notch1 with resistance to a PI3K inhibitor in T-cell leukaemia. Nature 513, 512–516 (2014). 10.1038/nature13495

48 Huang, B. J. et al. Convergent genetic aberrations in murine and human T lineage acute lymphoblastic leukemias. PLoS Genet 15, e1008168 (2019). 10.1371/journal.pgen.1008168

