## Supplementary Figures for "Acute Myeloid Leukemia Relapse after Bromodomain Inhibitor Treatment or Chemotherapy is Characterized by Myc-Ras Transcriptional Remodeling"

**Figure S1**

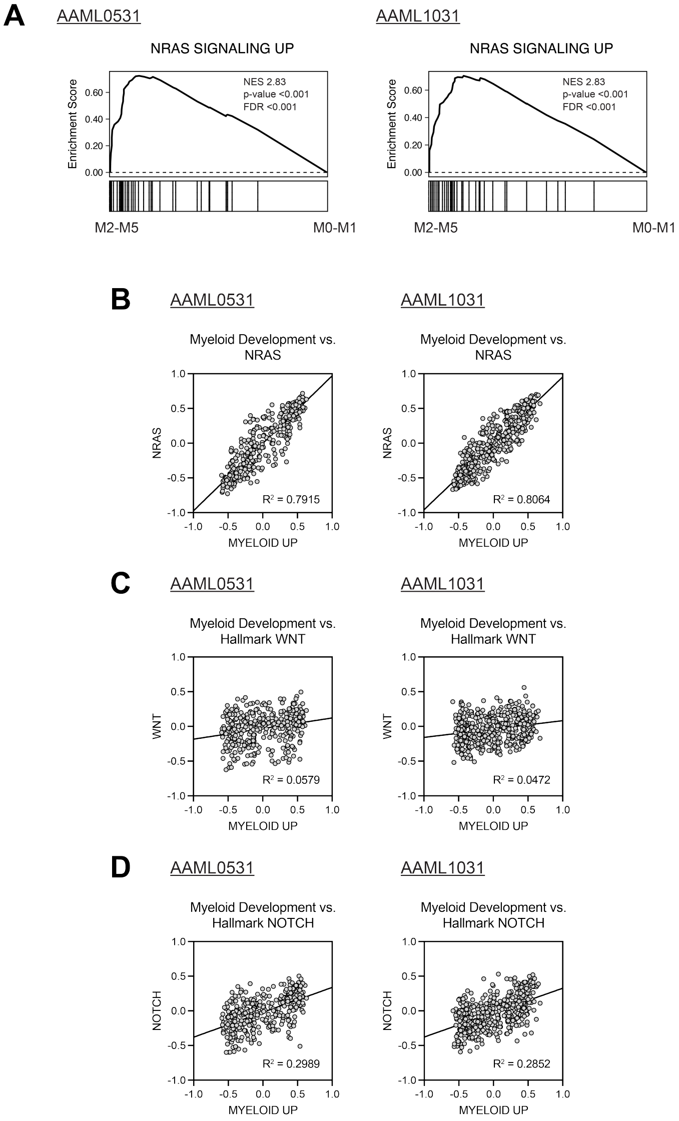

**Figure S1. Morphologic and transcriptional AML maturation is associated with increased Ras transcriptional output. A.** NCI TARGET AML bulk RNA sequencing (RNA-seq) data were analyzed in the context of French-American-British (FAB) AML morphologic classification. GSEA was performed across the Molecular Signature Database with a specific focus on Hallmark, C2 Curated, and C6 Oncologic Signature gene sets. The gene set Croonquist NRAS Signaling Up is enriched in FAB M2, M4, M5 AML subtypes (compared to FAB M0 and M1). **B.** GSVA enrichment scores for Broad Molecular Signature Database myeloid development versus NRAS signature gene sets are highly correlated in AAML0531 and AAML1031 patient cohorts. **C, D.** By contrast, lower $R^2$ values are observed with WNT and NOTCH gene sets.

### Figure S2

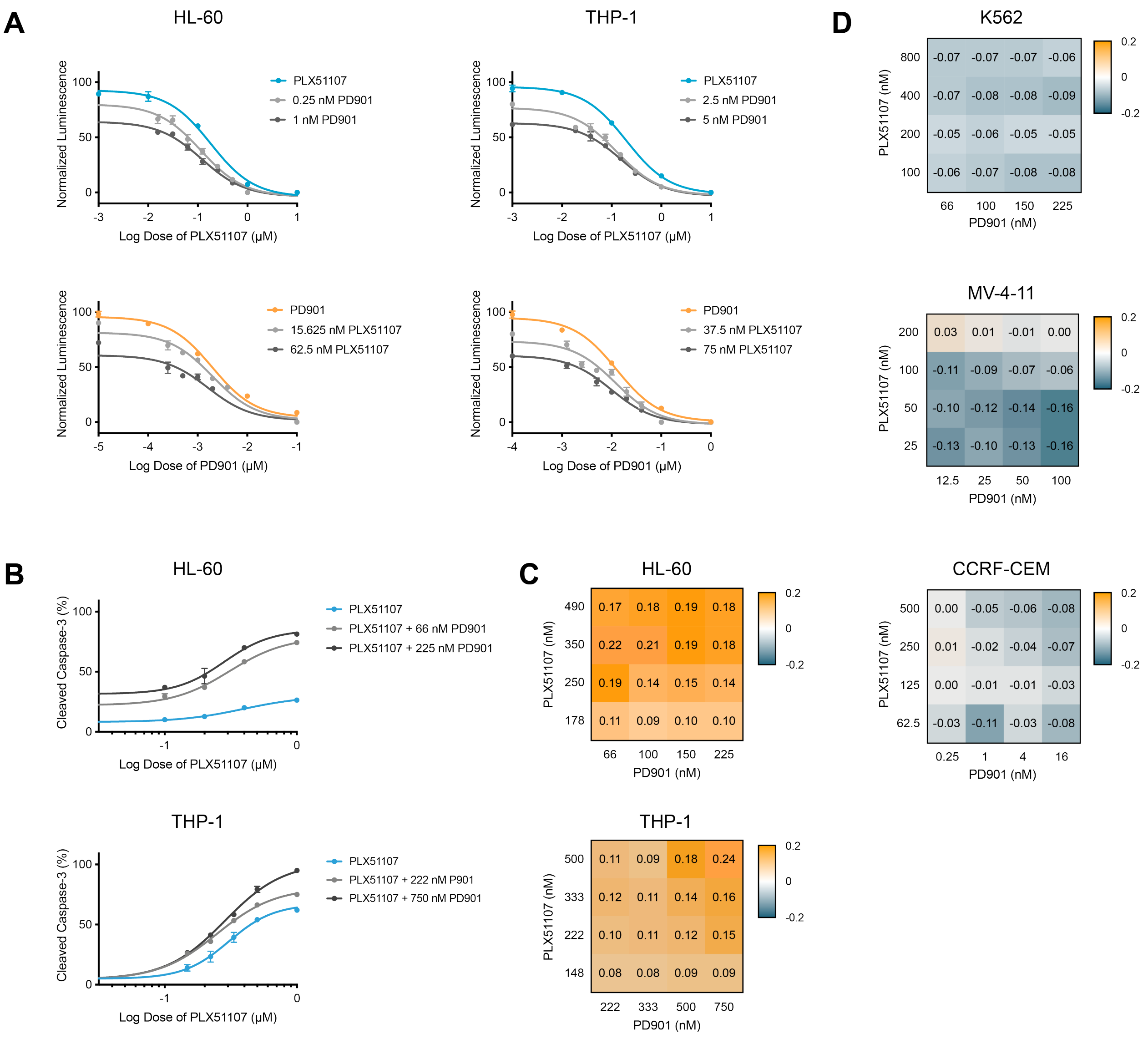

**Figure S2. Activity of PLX51107 and PD901 in NRAS-mutant monocytic AML cell lines. A.** CellTiter-Glo (CTG) analysis of the NRAS-mutant HL-60 and THP-1. **B, C.** Apoptosis as measured by the percentage of cells expressing cleaved-caspase-3 in response to PLX51107 ± PD901 in HL-60 and THP-1 cell lines. Positive (synergistic) Bliss independence scores were observed across a range of PLX51107 and PD901 doses. **D.** Negative (antagonistic) Bliss independence scores were observed in dose response experiments testing PLX51107 and PD901 in "negative control" cell lines (either not AML and/or not RAS mutant).

#### Figure S3

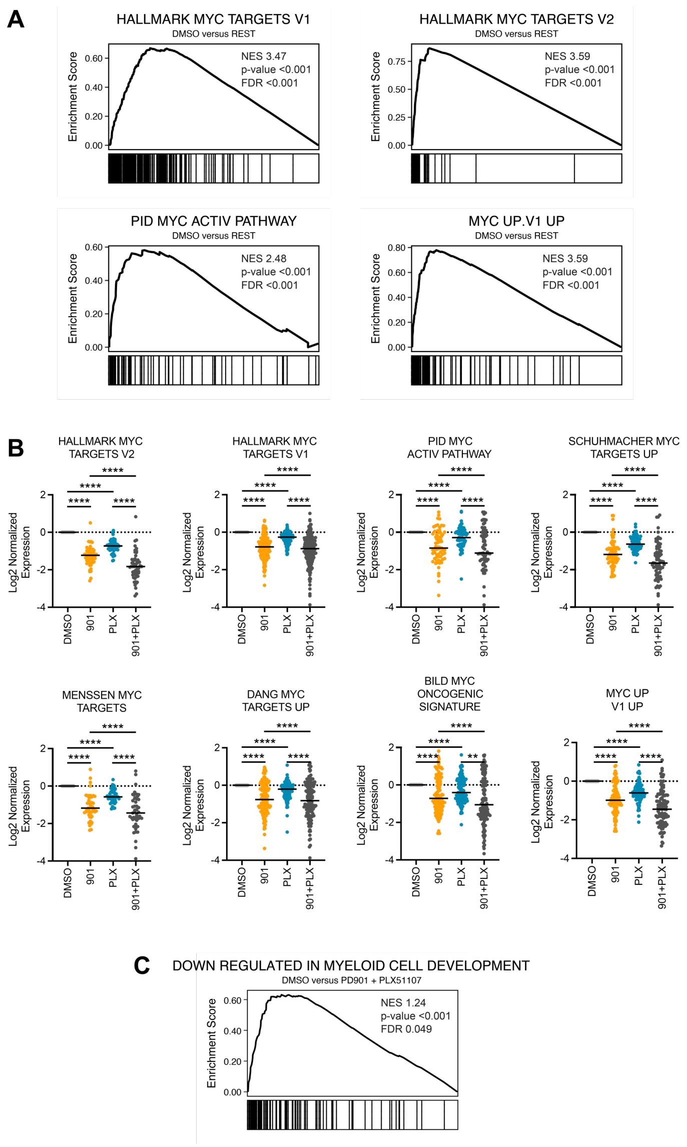

**Figure S3. PLX51107 and PD901 induce synergistic down-regulation of Myc regulated gene expression. A.** Bulk RNA sequencing (RNA-seq) analysis of OCI-AML3 cells treated with DMSO, PD901, PLX51107, or the combination. Myc regulated gene sets are enriched in DMSO treated cells compared to the other three drug treated conditions. **B.** Across a wide range of Myc regulated gene sets, the combination of PD901 and PLX51107 more potently down-regulates target genes than either agent alone. **C.** Treatment with PD901 and PLX51107 is induces a more immature myeloid transcriptional state.

#### Figure S4

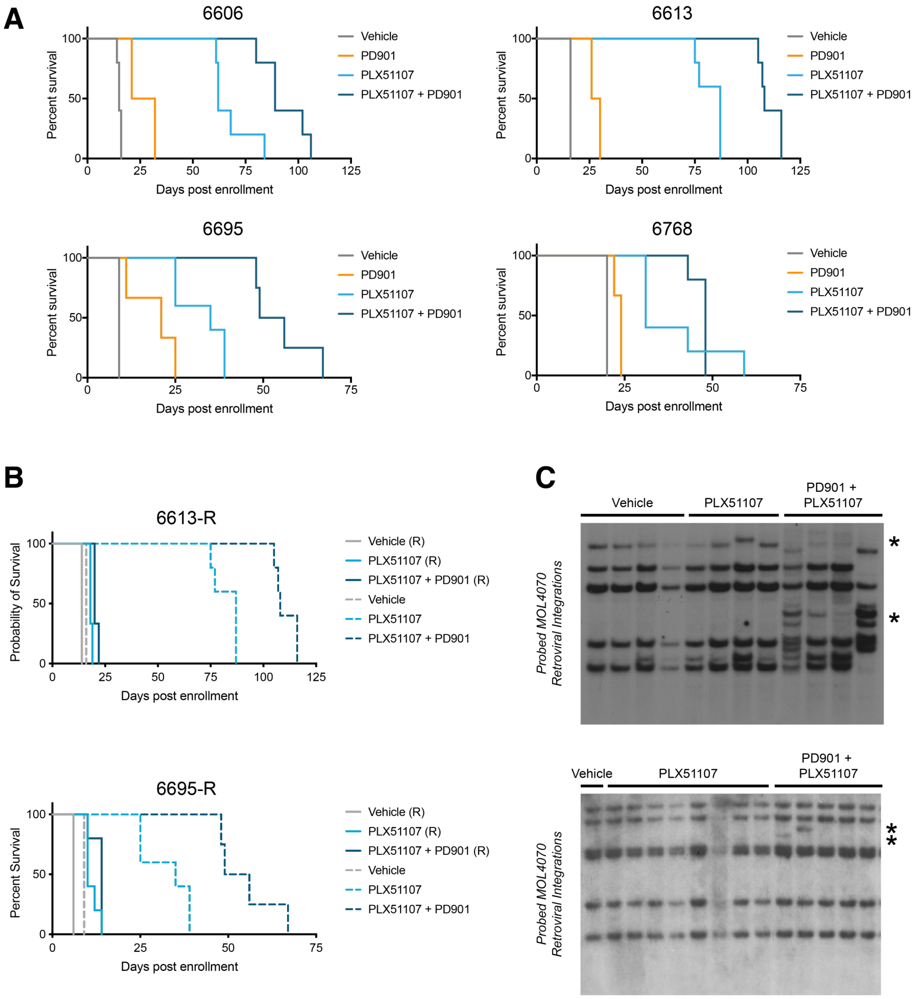

**Figure S4. Response and resistance of *Nras*$^{G12D}$ AMLs to PLX51107 and PD901. A.** Kaplan-Meier analysis of individual AMLs (6606, 6613, 6695, and 6768), which are shown in aggregate within Fig. 4A. **B.** Relapsed AMLs from the 6613 and 6695 were harvested from drug-treated moribund mice, retransplanted, and retreated with vehicle or the drug regimen used in the preceding primary preclinical trial (PLX51107 or PLX51107 + PD901). The in vivo sensitivity of retransplanted and retreated AMLs (solid line) is significantly reduced relative to the primary trial (dotted line). **C.** Southern blot probing for MOL4070LTR integrations was performed on DNA isolated from independent recipient mice transplanted with AML 6613 and 6695 that were treated with either vehicle, PLX51107, or PLX51107 + PD901. Novel restriction fragments are visible in mice treated with PLX51107 or PLX51107 + PD901 are indicated with asterisks.

#### Figure S5

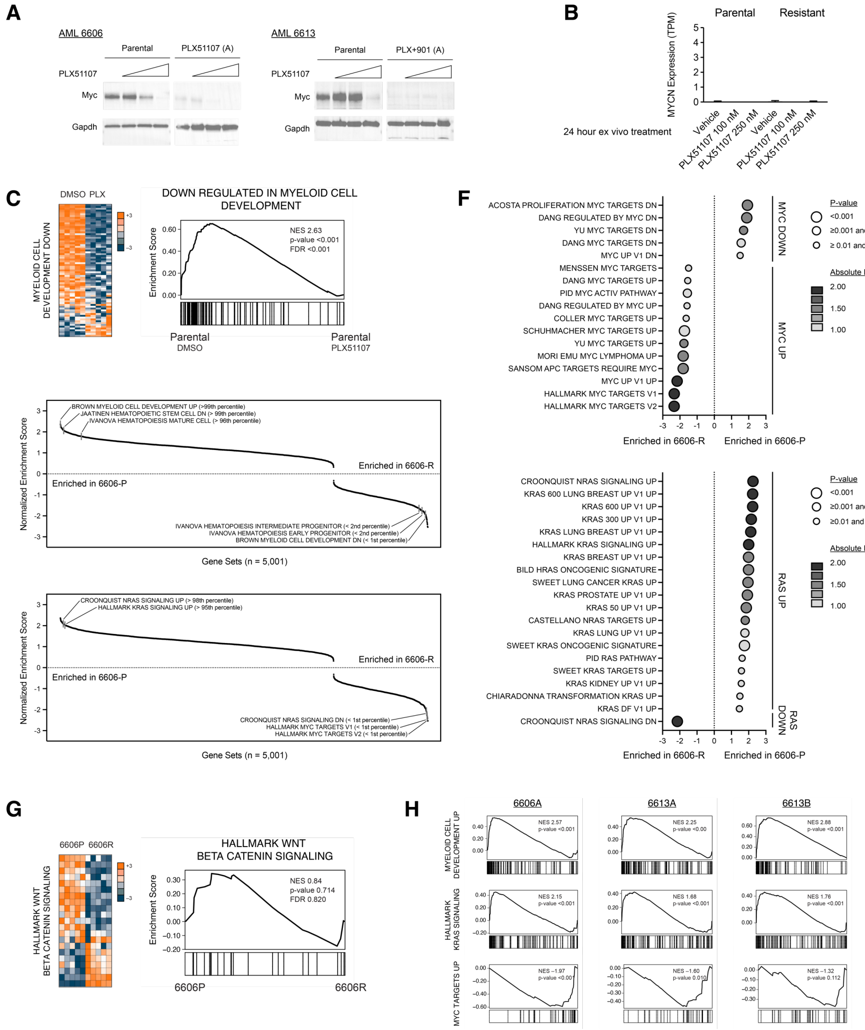

**Figure S5. Resistance to PLX51107 is associated with an inverse relationship between Myc and Ras transcriptional programs. A.** Exposing parental AML cells to PLX51107 ex vivo results in the expected down-regulation of Myc protein expression while confirmed resistant AML counterparts had either decreased or no detectable Myc protein. **B.** For AML 6606P and 6606B, no Mycn expression was detected either at baseline or in response to PLX51107. **C.** Treatment of AML 6606P *ex vivo* with PLX51107 250 nM for 24 hours induces a more immature myeloid transcriptional state. **D, E, F.** Bulk RNA sequencing was performed on AML6606P and 6606B, which revealed broad up regulation of Myc associated transcriptional programs and down-regulation of Ras programs in AML 6606B (compared to 6606P). **G.** However, Wnt/β-catenin signatures were not enriched. **H.** GSEA analysis of 6606A and 6613A/6613B (compared to 6606P and 6613P, respectively).

#### Figure S6

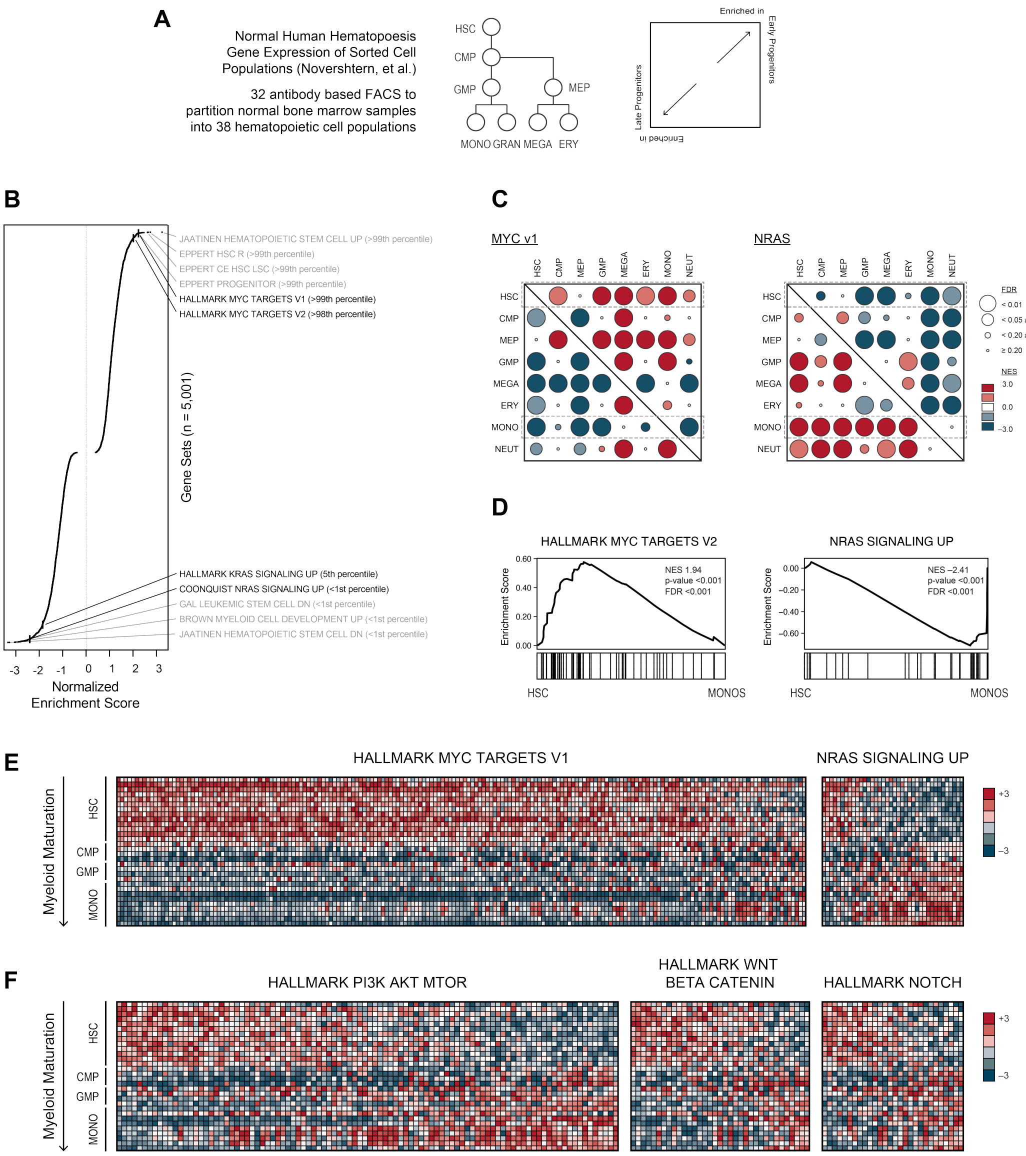

**Figure S6. Normal hematopoietic differentiation is associated with downregulated Myc and upregulated Ras programs. A.** Analysis performed on the normal hematopoiesis gene expression dataset published by Novershtern, et al. **B.** Comparing HSCs versus monocyte populations, GSEA was performed across the Broad Molecular Signature Database (Hallmark, C2, and C6 gene sets) with enrichment in hematopoiesis and Myc/Ras gene sets. **C.** Pair-wise GSEA comparisons between hematopoietic cell populations with a specific focus on Myc and Ras gene sets. **D.** GSEA plots, normalized enrichment scores, and false discovery rates. **E, F.** Ranked gene expression of MYC/RAS signatures versus other pathways (PI3K, WNT, and NOTCH), showing the former is associated with HSC/myeloid differentiation.

**Figure S7**

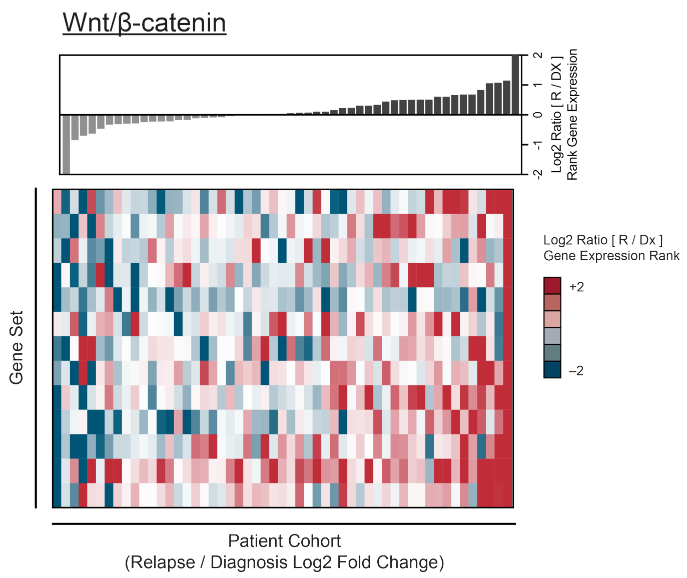

**Figure S7. Relapsed pediatric AMLs are not enriched for Wnt/β-catenin associated transcriptional programs.** $Log_2$ ratio gene expression fold change from diagnosis to relapse across the Hallmark Wnt/β-catenin signature gene set.

### Figure S8 (panel 1 of 2)

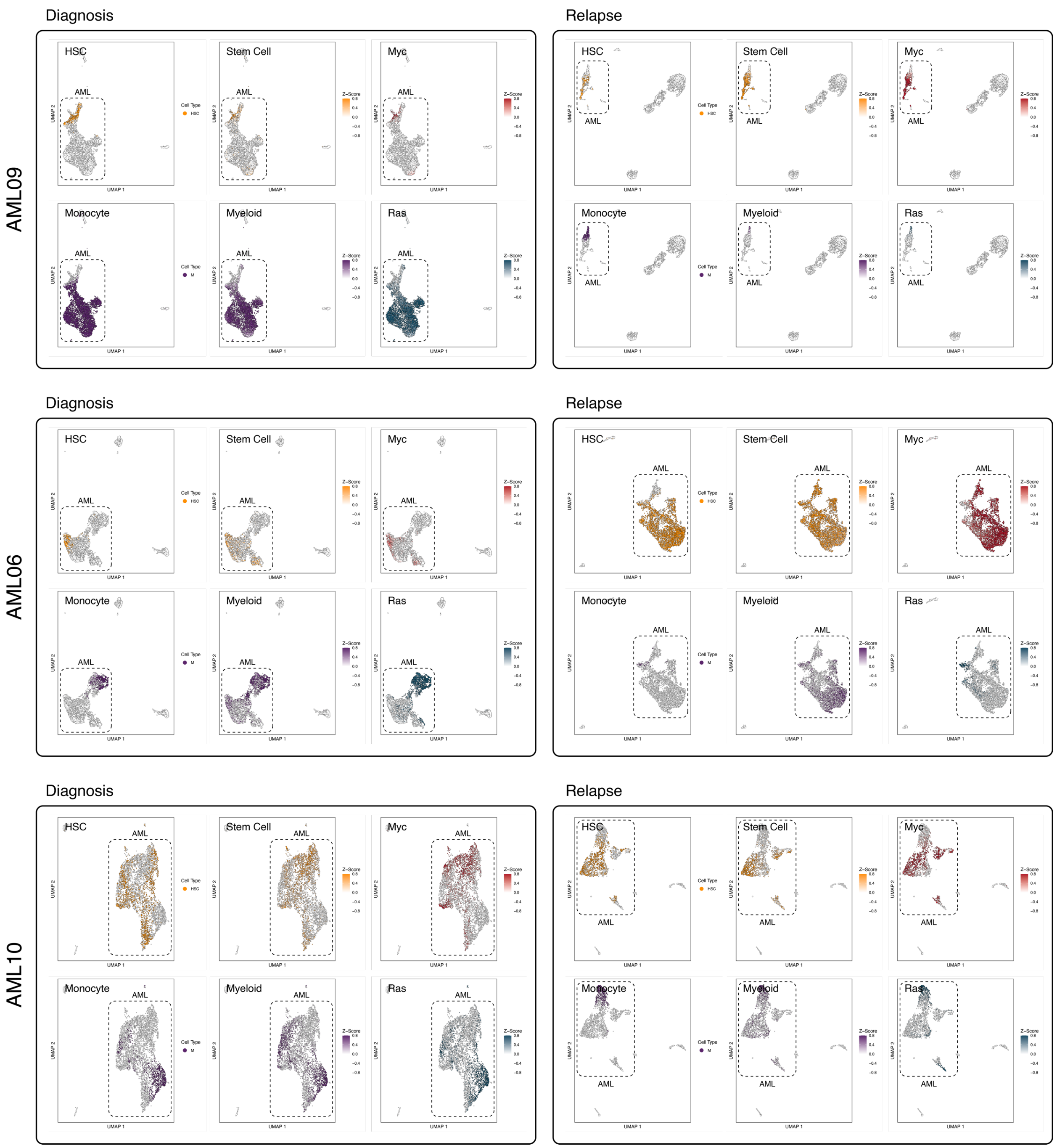

#### Figure S8 (panel 2 of 2)

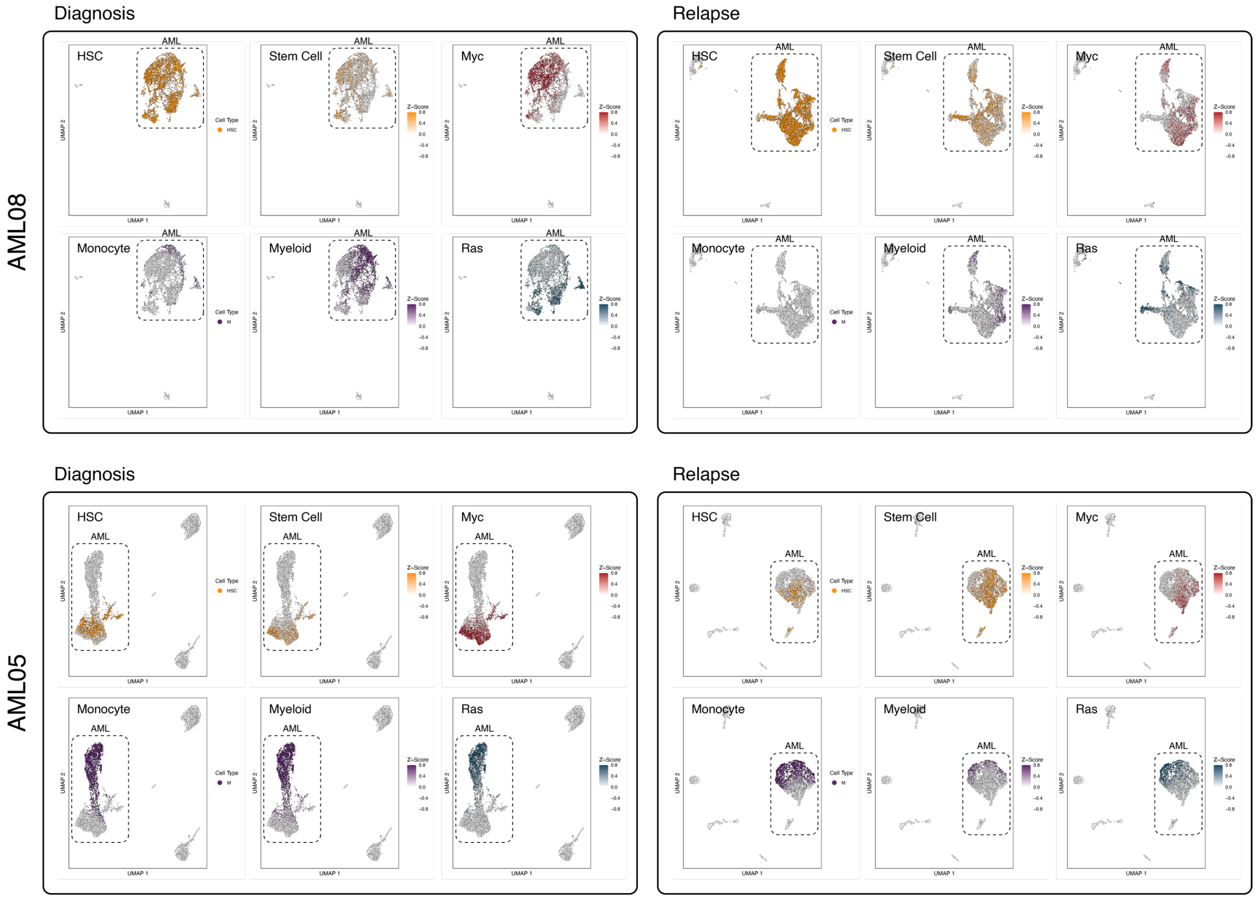

**Figure S8. Single cell RNA and ATAC sequencing show analogous patterns of transcriptional plasticity.** Single cell RNA and ATAC sequencing (scRNA/ATAC-seq) of diagnosis and relapsed KMT2A-rearranged AMLs reveals transcriptional transitions. HSC and Monocyte UMAP plots were assigned a color based on the assigned hematopoiesis maturation as described in Lambo, et al. Stem Cell, Myeloid, Myc, and Ras UMAP plots were shaded based on Broad Molecular Signature Database GSVA enrichment scores.

#### Table S1

| | | Non-Myeloid Cell Lines | | | Myeloid Cell Lines | | |
|---|---|---|---|---|---|---|---|
| Cell Line | Subtype | Mean | 95% CI | 95% CI | Mean | 95% CI | 95% CI |
| CCRF-CEM | Leukemia, acute lymphoblastic | 0.020 | 0.027 | 0.015 | | | |
| RPMI 8226 | Myeloma, plasmacytoma | 0.020 | 0.027 | 0.015 | | | |
| MV4-11 | Leukemia, biphenotypic B myelomonocytic | 0.031 | 0.032 | 0.029 | | | |
| L_8E5 | Leukemia, acute lymphoblastic | 0.051 | 0.081 | 0.032 | | | |
| SC | Monocyte/Macrophage | | | | 0.056 | 0.088 | 0.036 |
| KE-37 | Leukemia, T cell | 0.067 | 0.083 | 0.054 | | | |
| HuT 78 | Lymphoma | 0.067 | 0.090 | 0.050 | | | |
| OCI-AML3 | Acute myeloid leukemia | | | | 0.059 | 0.063 | 0.055 |
| SKM-1 | Acute myeloid leukemia | | | | 0.072 | 0.077 | 0.067 |
| Loucy | Leukemia, acute lymphoblastic t(16;20) | 0.084 | 0.110 | 0.067 | | | |
| BDCM | Leukemia, acute myelogenous | 0.084 | 0.130 | 0.055 | | | |
| MOLM14 | Acute myeloid leukemia | 0.089 | 0.117 | 0.061 | | | |
| RL | Lymphoma, non-Hodgkin's | 0.091 | 0.120 | 0.071 | | | |
| J.RT3-T3.5 | Leukemia, acute T cell | 0.100 | 0.130 | 0.081 | | | |
| CA46 | Lymphoma, Burkitt's | 0.110 | 0.160 | 0.076 | | | |
| BC-1 | Lymphoma, EBV and KSHV positive | 0.110 | 0.190 | 0.068 | | | |
| DOHH2 | Lymphoma | 0.120 | 0.140 | 0.096 | | | |
| EOL-1 | Lymphoma | 0.120 | 0.160 | 0.092 | | | |
| THP-1 | Acute myeloid leukemia | | | | 0.137 | 0.143 | 0.132 |
| CEM/C2 | Leukemia, acute lymphoblastic | 0.140 | 0.180 | 0.110 | | | |
| MV-4-11 | Acute myeloid leukemia | | | | 0.141 | 0.188 | 0.094 |
| K562 | Leukemia, myelogenous | | | | 0.150 | 0.230 | 0.100 |
| HL-60 | Acute myeloid leukemia | | | | 0.151 | 0.181 | 0.120 |
| MOLT-4 | Leukemia, acute lymphoblastic | 0.160 | 0.270 | 0.089 | | | |
| P3HR-1 | Lymphoma, Burkitt's | 0.160 | 0.210 | 0.130 | | | |
| Kasumi-1 | Leukemia, acute myeloblastic | | | | 0.180 | 0.230 | 0.140 |
| MPC-11 | Myeloma, Mouse | 0.190 | 0.260 | 0.140 | | | |
| H9 | Lymphoma, cutaneous | 0.220 | 0.240 | 0.200 | | | |
| Mo | Leukemia, hairy cell | 0.230 | 0.320 | 0.160 | | | |
| CEM/C3 | Leukemia, acute lymphoblastic | 0.240 | 0.310 | 0.180 | | | |
| MJ | Lymphoma, cutaneous T cell, mycosis fungoides | 0.240 | 0.320 | 0.180 | | | |
| JVM-2 | Leukemia | 0.240 | 0.400 | 0.150 | | | |
| HH | Lymphoma, cutaneous T cell | 0.250 | 0.390 | 0.150 | | | |
| NB-4 | Acute myeloid leukemia | | | | 0.256 | 0.264 | 0.247 |
| TF-1 | Erythroleukemia | 0.260 | 0.430 | 0.160 | | | |
| KU812 | Leukemia, chronic myelogenous | | | | 0.270 | 0.360 | 0.190 |
| NOMO1 | Acute myeloid leukemia | | | | 0.318 | 0.347 | 0.288 |
| AHH-1 | Lymphoblastoid | 0.300 | 0.410 | 0.220 | | | |
| GDM-1 | Leukemia, myelomonoblastic | | | | 0.340 | 0.480 | 0.230 |
| MOLT-3 | Leukemia, acute lymphoblastic | 0.390 | 0.730 | 0.210 | | | |
| KG-1 | Leukemia, acute lymphoblastic | 0.390 | 0.700 | 0.220 | | | |
| RPMI 7666 | Lymphoblast | 0.430 | 0.740 | 0.250 | | | |
| Daudi | Lymphoma, Burkitt's | 0.450 | 0.540 | 0.370 | | | |
| CESS | Leukemia, myelomonocytic | | | | 0.490 | 0.840 | 0.290 |
| GK-5 | B lymphoblast; Epstein-Barr virus (EBV) transformed | 0.530 | 0.910 | 0.300 | | | |
| ARH-77 | Leukemia, plasma cell | 0.570 | 1.200 | 0.270 | | | |
| CEM/C1 | Leukemia, acute lymphoblastic | 0.580 | 0.840 | 0.390 | | | |
| D1.1 | Leukemia, acute T cell, CD4 negative | 0.620 | 1.520 | 0.250 | | | |
| P116 | Leukemia, acute T cell | 0.620 | 1.030 | 0.370 | | | |
| SUP-B15 | Leukemia, acute lymphoblastic | 0.630 | 1.040 | 0.380 | | | |
| Jurkat | Leukemia, acute T cell | 0.670 | 1.080 | 0.420 | | | |
| KU 812 E | Leukemia, chronic myelogenous | | | | 0.730 | 0.840 | 0.630 |
| CCRF-HSB-2 | Leukemia, acute lymphoblastic | 0.820 | 1.280 | 0.520 | | | |

| | | | | | | | |
|---|---|---|---|---|---|---|---|
| WIL2-S | B lymphoblast, hereditary spherocytosis, spleen | 0.930 | 1.390 | 0.630 | | | |
| MC116 | Lymphoma, undifferentiated | 0.950 | 1.880 | 0.480 | | | |
| J45.01 | Leukemia, acute T cell, CD45 deficient | 0.990 | 1.400 | 0.700 | | | |
| P116.cl39 | Leukemia, acute T cell | 1.880 | 4.410 | 0.800 | | | |
| Ramos.2G6.4C10 | Lymphoma, Burkitt's | 1.910 | 3.830 | 0.960 | | | |
| RS4;11 | Leukemia, acute lymphoblastic, t(4;11) translocation | 2.130 | 6.160 | 0.730 | | | |
| SU-DHL-6 | Lymphoblast-like, peritoneal effusion | 2.860 | 9.460 | 0.860 | | | |
| I 9.2 | Leukemia, acute T cell | 2.880 | 8.770 | 0.940 | | | |
| TALL-1 | Lymphoma, T cell | 3.300 | 6.840 | 1.600 | | | |
| SUP-T1 | Leukemia, lymphoblastic | 3.700 | 8.560 | 1.600 | | | |
| Raji | Lymphoma, Burkitt's | 4.000 | 6.510 | 2.450 | | | |
| ST486 | Lymphoma, Burkitt's | 5.190 | 8.990 | 3.000 | | | |
| U266B1 | Myeloma; plasmacytoma | 6.590 | 12.200 | 3.560 | | | |
| U-937 | Lymphoma, histiocytic | 7.580 | 10.620 | 5.420 | | | |
| AML-193 | Leukemia, Acute monocytic | | | | 8.210 | 13.940 | 4.840 |
| Toledo | Lymphoma, diffuse large cell, non-Hodgkin's B cell | 12.060 | 64.510 | 2.250 | | | |
| NAMALWA | Lymphoblastoid | 13.570 | 19.440 | 9.470 | | | |
| EB-1 | Lymphoma, Burkitt's | >20 | | | | | |
| EB-2 | Lymphoma, Burkitt's | >20 | | | | | |
| GA-10 | Lymphoma, Burkitt's | >20 | | | | | |
| NCI-H929 | Myeloma, plasmacytoma | >20 | | | | | |

#### Table S2

| AML | Gene Symbol(s) | Treatment Arm (Clone #)<br>Total Insertions (%) | | |
|---|---|---|---|---|
| 6606 | | Vehicle (40) | PLX51107 (20) | PLX51107 + 901 (16) |
| | Alg13 \| Trpc5 | 16 (40) | 14 (17) | 2 (2) |
| | Sox4 | 15 (30) | 30 (37) | 19 (18) |
| | Cd72 \| Sit1 \| Tesk1 \| Rmrp | 9 (17) | 12 (15) | 12 (12) |
| | Mir23a \| Mir27a \| Mir24-2 | – | 12 (15) | – |
| | Nol4l \| Commd7 \| Dnmt3b | 1 (2) | 3 (4) | 45 (43) |
| 6613 | | Vehicle (0) | PLX51107 + PD901 (26) | PLX51107 + 901 (30) |
| | Rad51b | 34 (22) | 14 (8) | 25 (17) |
| | Dusp22 \| Irf4 | 30 (19) | 13 (10) | 30 (20) |
| | Mecom \| Mannr | 21 (13) | 8 (5) | 13 (9) |
| | Sox4 | 2 (1) | 27 (16) | – |
| | Bcl9l \| Cxcr5 \| Upk2 \| Foxr1 | – | – | 15 (10) |
| | Ptprf \| Mir7226 \| Kdm4a | – | – | 13 (9) |
| | Myb | – | – | 2 (1) |
| 6695 | | Vehicle (12) | PLX51107 + 901 (30) | PLX51107 + 901 (32) |
| | Sh2d4a | 25 (24) | 29 (30) | 26 (29) |
| | Pcyox1l \| Grpel2 \| Il17b | 24 (23) | 14 (15) | 12 (14) |
| | Sox4 | 23 (22) | 6 (6) | 9 (10) |
| | Mir3471-2 \| Tmem68 \| Tgs1 | – | 4 (4) | 3 (3) |
